# Genetic and clinical analyses of psychosis spectrum symptoms in a large multi-ethnic youth cohort reveal significant link with ADHD

**DOI:** 10.1101/814087

**Authors:** Loes Olde Loohuis, Eva Mennigen, Anil Ori, Diana Perkins, Elise Robinson, Jean Addington, Kristin S. Cadenhead, Barbara A. Cornblatt, Daniel H. Mathalon, Thomas H. McGlashan, Larry J. Seidman, Matcheri Keshavan, William Stone, Ming T. Tsuang, Elaine F. Walker, Scott W. Woods, Tyrone D. Cannon, Ruben C. Gur, Raquel E. Gur, Carrie E. Bearden, Roel A. Ophoff

**Affiliations:** Center for Neurobehavioral Genetics, Semel Institute for Neuroscience and Human Behavior, University of California, Los Angeles, California, USA; Department of Psychiatry and Psychotherapy, University Hospital Carl Gustav Carus, Technische Universität Dresden, Dresden, Germany; Department of Psychiatry, University of North Carolina, Chapel Hill, NC, USA; Stanley Center for Psychiatric Research, Broad Institute of MIT and Harvard, Cambridge, Massachusetts, USA; Program in Medical and Population Genetics, Broad Institute of MIT and Harvard, Cambridge, Massachusetts, USA; Department of Epidemiology, Harvard T.H. Chan School of Public Health, Boston, Massachusetts, USA; Department of Psychiatry, Hotchkiss Brain Institute, Calgary, Alberta, Canada; Department of Psychiatry, UCSD, San Diego, CA; Department of Psychiatry, Zucker Hillside Hospital, Long Island NY; Department of Psychiatry, UCSF, and SFVA Medical Center, San Francisco CA; Department of Psychiatry, Yale University, New Haven CT; Department of Psychiatry, Harvard Medical School at Beth Israel Deaconess Medical Center and Massachusetts General Hospital, Boston MA; Departments of Psychology and Psychiatry, Emory University, Atlanta GA; Department of Psychology, Yale University, New Haven CT; Department of Psychiatry, University of Pennsylvania School of Medicine and the Penn-CHOP Lifespan Brain Institute; Philadelphia, USA; Department of Psychology, University of California, Los Angeles, California, USA; Department of Human Genetics, David Geffen School of Medicine, University of California, Los Angeles, California, USA; Department of Psychiatry, Erasmus University Medical Center, Rotterdam, The Netherlands

## Abstract

**Objective:** Psychotic symptoms are an important feature of severe neuropsychiatric disorders, but are also common in the general population, especially in youth. The genetic etiology of psychosis symptoms in youth remains poorly understood. To characterize genetic risk for psychosis spectrum symptoms (PS), we leverage a community-based multi-ethnic sample of children and adolescents aged 8-22 years, the Philadelphia Neurodevelopmental Cohort (n = 7,225, 20% PS).

**Methods:** Using an elastic net regression model, we aim to classify PS status using polygenic scores (PGS) based on a range of heritable psychiatric and brain-related traits in a multi-PGS model. We also perform univariate PGS associations and evaluate age-specific effects.

**Results:** The multi-PGS analyses do not improve prediction of PS status over univariate models, but reveal that the attention deficit hyperactivity disorder (ADHD) PGS is robustly and uniquely associated with PS (OR 1.12 (1.05, 1.18) P = 0.0003). This association is: i) driven by subjects of European ancestry (OR=1.23 (1.14, 1.34), P=4.15×10^−7^) but is not observed in African American subjects (P=0.65) and ii) independent of phenotypic overlap. We also find a significant interaction with age (P=0.01), with a stronger association in younger children. In an independent sample, we replicate an increased ADHD PGS in 328 youth at clinical high risk for psychosis, compared to 216 unaffected controls (OR 1.06, CI(1.01, 1.11), P= 0.02).

**Conclusions:** Our findings suggest that PS in youth may reflect a different genetic etiology than psychotic symptoms in adulthood, one more akin to ADHD, and shed light on how genetic risk can be investigated across early disease trajectories.

## Introduction

Psychotic symptoms, such as delusions and hallucinations, are an important feature of severe psychiatric disorders such as schizophrenia and bipolar disorder. They are, however, also common in the general population, and occur in ∼5-10%(1) of adults; a prevalence much higher than that of clinical diagnoses of schizophrenia and bipolar disorder (about 0.5-3% each(2, 3)) In children and adolescents, the prevalence of psychotic symptoms/psychotic-like experiences is even higher, as high as 20%(4). Youth experiencing psychotic symptoms typically exhibit a multitude of other comorbid symptoms, such as increased mood, anxiety, and attention deficit hyperactivity disorder (ADHD) symptoms, as well as increased substance use and impairments in global functioning(5-7). While subclinical psychopathology poses a risk for later development of overt psychiatric illnesses(5, 8-11), only a minority of youth reporting psychotic symptoms will convert to full-blown psychotic disorders.

With recent progress in psychiatric genetics, psychotic disorders are becoming well-characterized genetically(12-14). In particular, the landmark genome-wide association study (GWAS) of schizophrenia provides aggregate risk conferred by variants identified, polygenic scores (PGS), which explain about 7-10% of variance in case-control status(12, 15). In individuals with bipolar disorder, both genetic risk for schizophrenia as well as for bipolar disorder have been associated with psychotic symptoms(16, 17).

In the general population, however, the genetic etiology of psychotic symptoms across development is still largely unknown. The heritability of psychotic experiences has been estimated between 30-50% from twin studies(18, 19) with the proportion of genetic variance explained by common variants (SNP-heritability) of 3-17% in adolescents(18, 20). Adults with psychotic symptoms harbor increased genetic liability for a broad spectrum of psychiatric disorders, including schizophrenia and other neuropsychiatric disorders(21). While in adolescents some evidence suggests increased genetic risk for schizophrenia (and major depressive disorder) for specific features of psychosis(18), the reported effect sizes are very small, and these effects not very robust(22). In pre-adolescent youth, the relationship between genetic risk for psychiatric traits and psychotic symptoms has not yet been explored. The genetic characterization of psychosis spectrum symptoms across development may increase our understanding of their etiological and pathological significance.

To study genetic risk for psychosis symptoms in a population sample of youth, we leverage a large well-characterized community-based sample of youth aged 8-22 years, the Philadelphia Neurodevelopmental Cohort (PNC, n = 9,498 in total). The PNC is a multi-ethnic cohort, with the largest proportion of individuals of European (66%) and African American (26%) ancestry. In this community-based sample that is not ascertained for neuropsychiatric disorders, about 20% of the youth are classified as having psychotic spectrum symptoms (PS). In the PNC, having PS has been associated with structural and functional brain alterations(23-25), qualitatively similar to those present in overt psychotic disorders, as well as cognitive deficits(26). These findings underscore both the increased vulnerability in youth experiencing psychosis spectrum symptoms and the importance of investigating psychosis risk as a dynamic developmental process(7).

In this well-characterized sample, we explore the genetic architecture of PS based on common variant liability for psychiatric illness. Specifically, we aim to understand whether PGS for psychiatric disorders can be used to classify psychotic symptoms in youth. To do so, we adopt a recently developed multi-PGS approach (27). The method combines multiple summary statistics from different GWAS into a single predictive model, thus potentially increasing classification power. Given evidence suggesting PS increases risk for broader psychopathology(11, 21) and the wide spectrum of genetic inter-correlations in adults with psychotic experiences(21), we include a range of heritable psychiatric and brain traits in our analyses, an analytic approach that has not been previously explored in this type of cohort. In addition to multi-variate classification, we perform univariate associations and also assess phenotypic overlap between traits.

We hypothesize that developmental changes in the expression of psychotic symptoms will be reflected via age-specific genetic etiology, with the genetic architecture underlying psychosis spectrum symptoms in older youth being more similar to psychotic disorders. We evaluate this hypothesis by testing whether the observed correlations and the interplay with phenotypic overlap changes across the development.

## Methods

### Cohort Description

Data were obtained from the publicly available Philadelphia Neurodevelopmental Cohort (PNC, 1st release, phs000607.v1.p1, #7147) via the Database of Genotypes and Phenotypes (dbGap) platform. The PNC is a community-based sample consisting of 9,498 genotyped youth (ages 8-22 years) who participated in clinical and neurocognitive assessment, with a subsample undergoing MRI, after providing written informed consent or assent with parental consent (youth under 18 years old). Psychiatric symptomatology was assessed using the GOASSESS interview(28) covering broad domains of psychopathology including mood, anxiety, phobias, psychosis and externalizing behavior(29). Psychotic symptoms were specifically assessed with questions from the Kiddie Schedule for Affective Disorders and Schizophrenia for School-Age Children (K-SADS)(30), the Structured Interview for Prodromal Syndromes (SIPS)(31), and the PRIME Screen Revised (P-SR).

### Definition of psychosis spectrum

Criteria to establish a group of individuals that experiences psychosis spectrum symptoms were defined as in prior PNC studies(7, 24, 32). These criteria consider lifetime occurrence of positive psychotic symptoms such as hallucinations and persecutory thinking, negative/disorganized symptoms such as flattened affect, as well as age-appropriateness of these symptoms (Supplementary Methods). Previous publications have described the clinical and functional significance of these criteria(7, 24, 32).

Since MRI data are available for only a subset of the PNC sample that is too small for genetic analyses, we do not include these in our analyses. Other symptoms, such as PS domain scores of positive (PRIME) and negative/disorganized (SOPS) as well as symptoms of ADHD are described in the Supplementary Material.

### Genotyping QC and imputation

Genotyping QC, imputation and selection of individuals of European and African ancestry are described in detail in the Supplementary Materials. In brief, imputation followed the standard Ricopili pipeline (see URLS) and best-guess genotypes of well-imputed variants (INFO>0.8) were selected for further analysis. The imputed dataset included 7,774 subjects, 7,764 with phenotyping data. Individuals were assigned ancestry group based on estimates from ADMIXTURE(33) and related subjects (specifically 477 siblings) were removed within ancestry groups (Supplementary Note). After these final filtering steps, the total sample size is 7,225, with a total of 1,937,561 included SNPs. We identified two sub-cohorts: one including individuals of European ancestry (EA, N=4,852) and one African American ancestry group (AA, N=1,802).

### GWAS summary statistics

We selected GWAS summary statistics from LD hub (34), a centralized repository for summary statistics (accessed June 2018). Specifically, we included 23 GWAS summary statistics of psychiatric, brain traits and personality traits, that were either publicly available or obtained via personal correspondence. From these, a total of 12 had a linkage disequilibrium (LD) score heritability z-score >5, indicating good statistical power (27)(which is a function of variance explained and sample size) and complete GWAS information available. If available, we replaced summary statistics in LD hub with more recent or powerful GWAS: For the Psychiatric Genomics Consortium (PGC), we replaced MDD, BIP, ADHD. Finally, we also included the 23andMe traits for Morningness and self-reported depression. For more details on the included GWAS see Supplementary Table 1.

After filtering, the 13 traits included in our analyses include psychiatric disorders (PGC GWAS for: ADHD(35), Autism(36), bipolar disorder(13), schizophrenia(12), cross-disorder, a joint analyses of severe mental illness (37), major depression(38); other psychiatric GWAS for: self-reported depression from 23andMe (39) major depressive disorder from CONVERGE(40)) brain traits (ENIGMA Caudate volume and Putamen volume(41)) and behavioral traits (Morningness 23andMe(42), Neuroticism(43) and subjective well-being(43)(SWB)).

### Polygenic scoring

Polygenic scoring was performed in a standard clumping and thresholding fashion, based on a p-value threshold of 0.05 (see Supplementary Methods for details). Specifically, for analyses only involving the EA (or AA) cohort, we included ancestry-specific principal components (PCs) after exclusion of related samples. The standardized residuals were used for follow-up analyses.

In addition to the conventional approach of thresholding and clumping, which can lead to loss of information, especially in cases where the ancestry of the GWAS sample diverges from the target sample, we also computed polygenic scores using LDPRED for follow-up analysis(44). As recommended, we used the target sample genotype data as the LD reference panel, performing scoring separately in the EA and AA ancestry samples. We used standard settings with an LD radius of 500 SNPs.

### Multi-score PGS analyses

As has been successfully implemented previously (27), we used elastic net regularized regression to predict outcomes by selecting predictors and estimating their contribution to the prediction. Elastic net uses a linear combination of two regularization techniques, L2 regularization (used in ridge regression) and L1 regularization (used in LASSO), and has been shown to work particularly well in case of correlated predictors, as is expected in the context of highly correlated genetic traits(45).

Elastic net regularized regression employs two hyper-parameters, alpha and lambda. To achieve optimized balance between variance explained and minimum bias, we fit models to tune over both alpha and lambda parameter values in repeated 10-fold cross-validation, and used the minimal lambda for the prediction model. As a performance measure, we use area under the ROC curve (AUC). Models were trained on a random subset of 70% of the data and weights of the selected variables subsequently used to test their cumulative discriminatory power in predicting psychosis status in the remaining 30%. We performed model fit both in the entire sample as well as within EA and AA separately. To obtain an estimate of the robustness and range of the selected parameters we performed 1000 repetitions of the procedure. In addition, we also generated models for 1000 permutations of the phenotype (within the All, EA and AA cohorts). Comparing our models to those derived from permutation, we adopt the conservative approach to compare the mean of the 1000 repetitions in the actual sample to the distribution of the permutations.

### Univariate PGS analyses

To estimate the effect of each PGS individually, we fit a series of logistic regression models for each of the corrected polygenic scores including Age, Age^2^ and Sex as covariates. Effect sizes are reported as odds ratio (OR) relative to one standard deviation increase in PGS. To account for multiple testing, we applied a Bonferroni correction, dividing the p-value by the total number of tests (14×3=42 tests yields a p-value cutoff of 0.001). Interactions were tested by including the interaction term in the full model.

### Analyses of phenotyping data

Symptom overlap was tested using Fisher’s exact test (pairs of binary traits), Wilcoxon Rank-sum test (binary vs quantitative) and Spearman rank correlation (pairs of quantitative traits).

## Results

### Ancestry

After quality control (see Supplementary methods), the PNC cohort consists of a total of N=7,225 youth ages 8-22 with both phenotypic and imputed data available with two ancestry groups: European ancestry (EA, N=4,852) and African American ancestry (AA, n=1,802). From the total sample, 1,369 youth (19%) are classified as having psychosis spectrum symptoms (PS). Figure 1 displays the demographics of the cohorts. Common variant heritability of the PS phenotype was estimated using GCTA, but due to lack of power, we were unable to obtain an accurate estimate (SNP-h^2^ = 0.11, se=0.21, P=0.3, Supplementary Methods).

**Figure 1.**
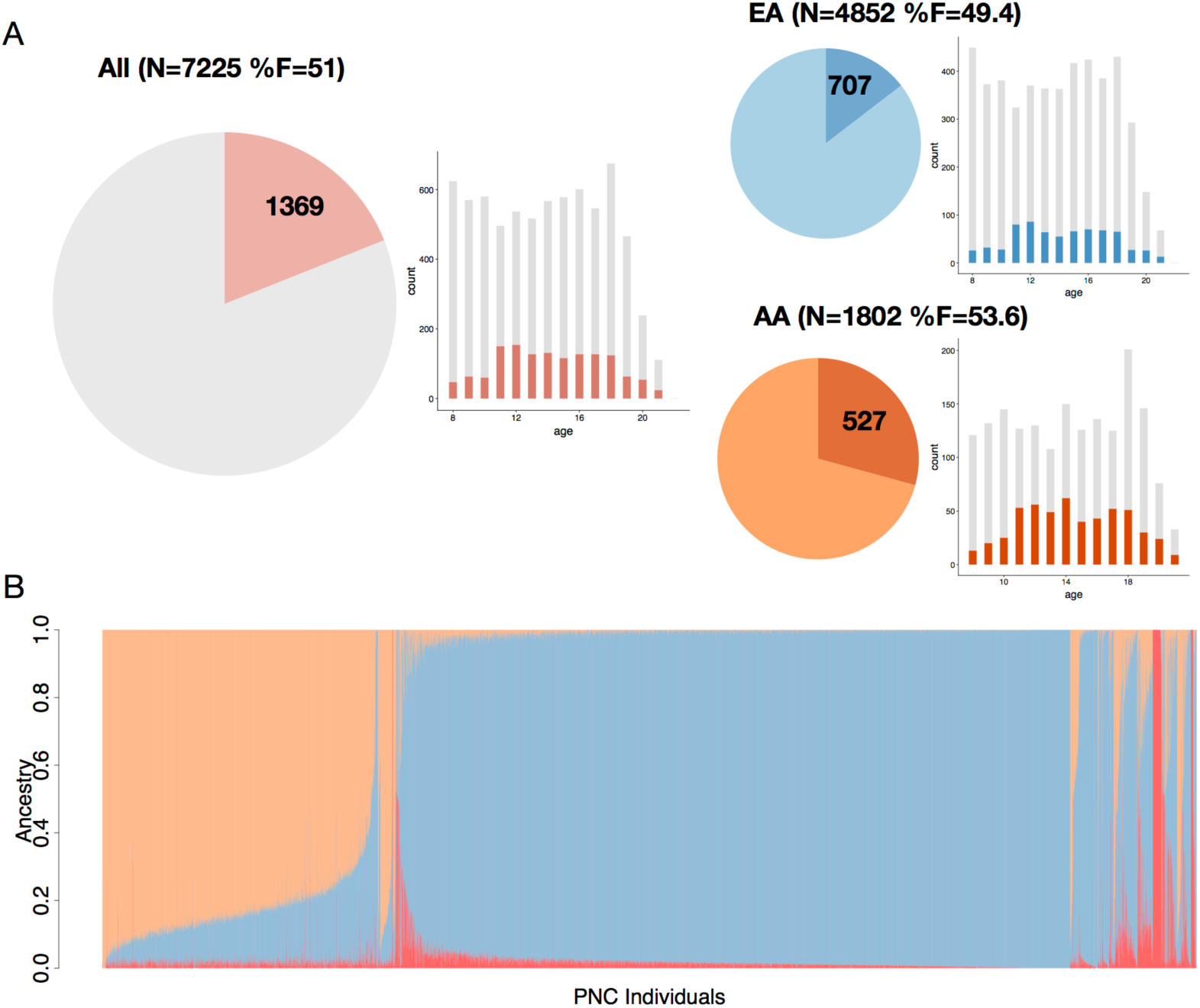
Demographics of the PNC cohort. 1a) Demographic overview of all subjects included in our analyses: pie charts display the proportion of subjects with psychosis spectrum symptoms (PS), and histograms the age distributions, with darker colors indicating PS subjects. Three groups included are all subjects (All), subjects of European and African American ancestry (EA and AA respectively). 1b) Relative admixture ancestry components (based on K=3) for the PNC cohort, ordered by self-reported ancestry.

### Multi-PGS prediction

Our multi-PGS models classified PS marginally better than chance, both in the whole sample and in the EA cohort alone (average AUC All =0.53 (sd 0.01) P=0.009, average AUC EA=0.55 (sd 0.02) P< 0.001). Within the AA cohort, however, the multivariate prediction was not different from chance (average AUC AA =0.51 (sd 0.02) P=0.35). In EA, the highest weight was consistently assigned to the PGS of ADHD, with an average standardized coefficient of 0.09 (sem 0.0007) corresponding to an OR of 1.10 (Figure 2), after correcting for all other selected variables. This effect was driven by the EA cohort (OR= 1.18; Figure 2 and Supplementary Figure 1) but absent in AA (OR=1.00). As expected, permutation of case-control status separately within the ancestry groups did not highlight any single trait (Supplementary Figure 4).

**Figure 2.**
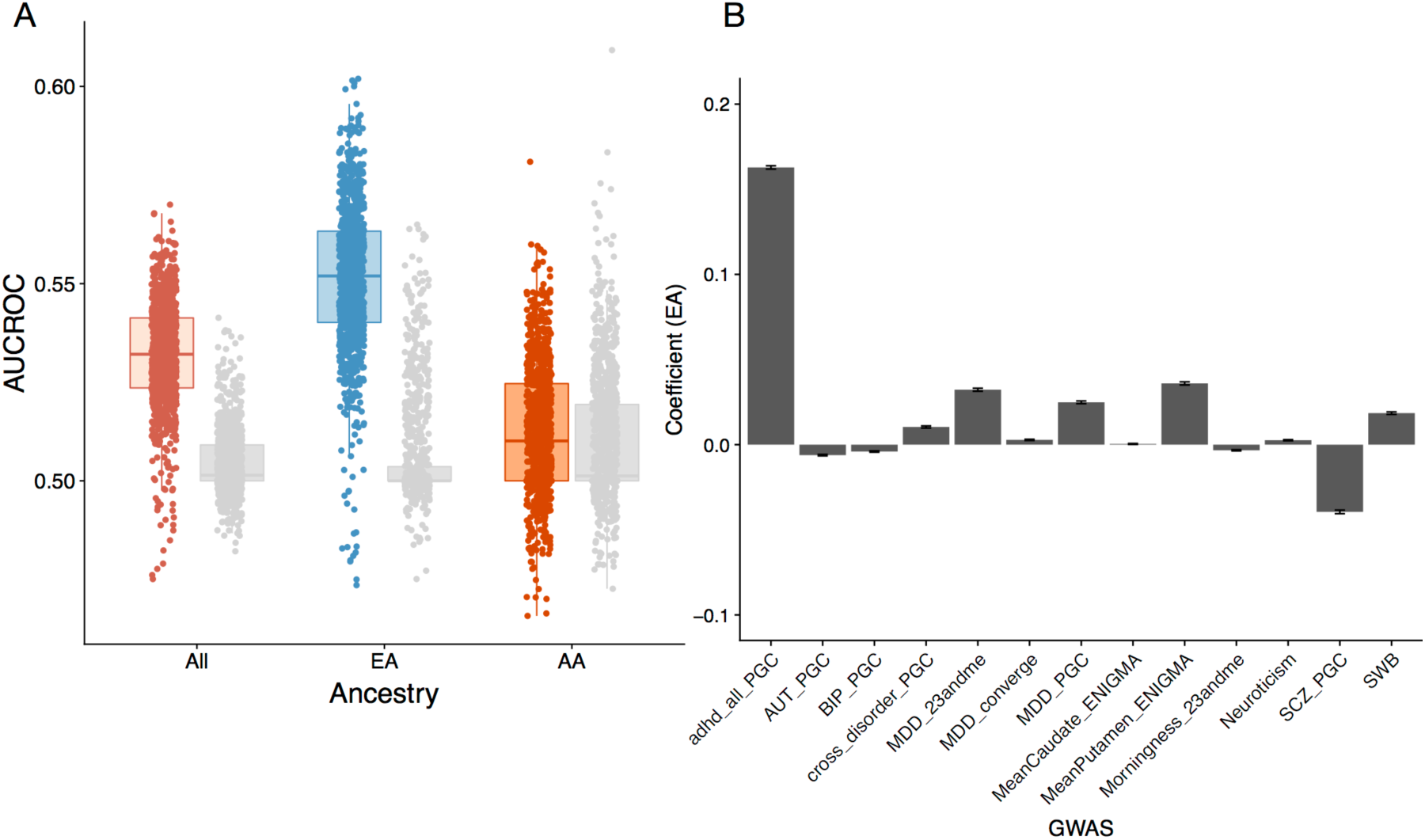
Multivariate classification of PS, by ancestry. 2a) AUCROC for each elastic net model trained on 70% and tested on the remaining 30% of data in All (pink), EA (blue) and AA (orange). Box plots indicate the median and the lower and upper hinges correspond to the first and third quartiles. The grey dots and boxplots refer to fits of permuted datasets within each ancestry group. The observed predictive power is driven by the EA cohort. 2b) Mean regression coefficients for the PGS based on different GWAS in EA. Standard errors indicate standard errors of the mean. The ADHD PGS is consistently included in the regression with the highest weight.

**Figure 3.**
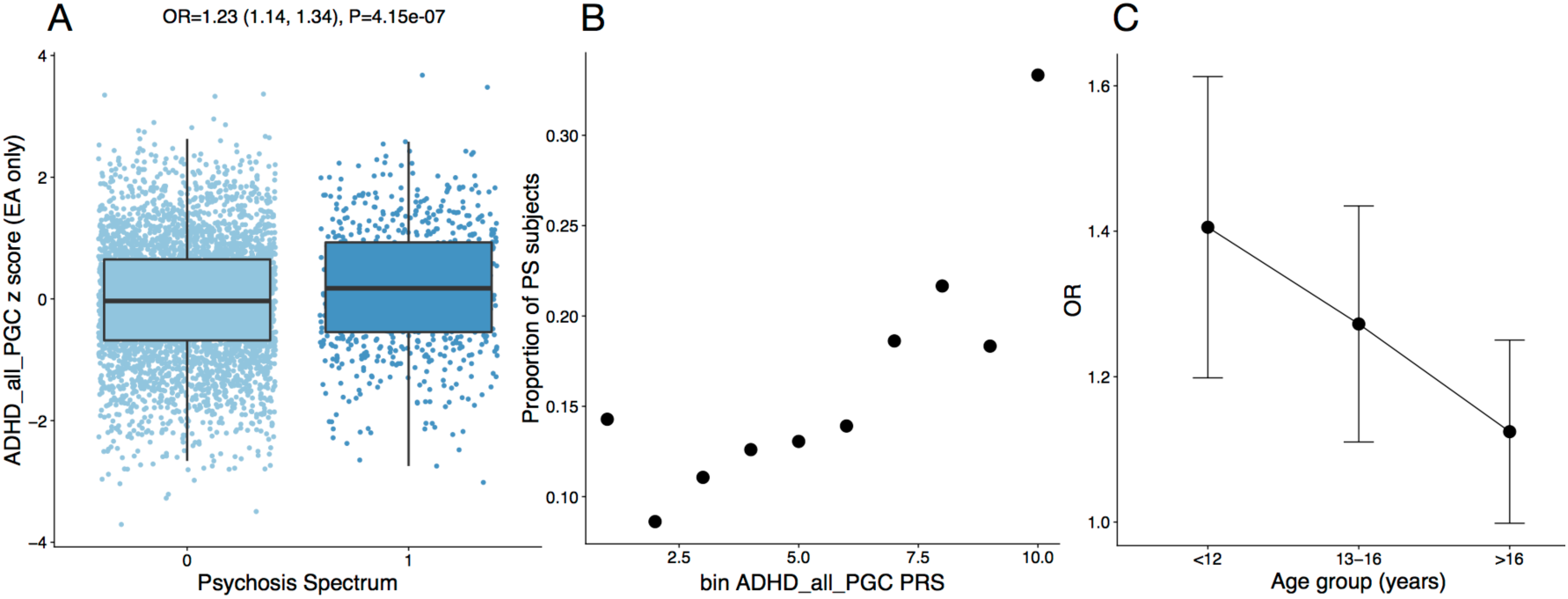
Univariate regression of ADHD PGS with PS in EA. 3a) Standardized ADHD PGS are higher in EA youth with PS versus non-PS youth. Each SD increase in PGS is associated with an OR=1.23 CI=(1.14, 1.34), P=4.15e-07. b) Proportion of cases with PS per PGS decile. c) Relative association in different age-bins (8-12, 12-16 and 16-22). The association between ADHD PGS and PS is strongest in youngest children 12 and younger. Adding the interaction term of age:PGS to the full model is significant (P=0.01).

### Univariate association

In line with the multivariate model, univariate logistic regression yielded a modest but significant association between PS and ADHD liability (OR 1.12 (1.05, 1.18) P = 0.0003, Figure 2, Table 1, Figure S6). This effect was driven by youth with EA ancestry (OR=1.23 (1.14, 1.34), P=4.15×10^−7^), and not observed in the AA cohort (OR=0.98 (0.88,1.08) P=0.65). Genetic liability for all other neuropsychiatric traits, including schizophrenia, was not associated with PS in either ancestry group. Since the AA cohort is of a different ancestry than the majority of GWAS cohorts, with different allele frequencies, LD patterns and effect sizes(46, 47), we also performed PGS computation using LDPRED(44), a method that explicitly models LD. No other traits were significantly associated using this method either. To test whether the multivariate predictor outperformed ADHD PGS alone, we performed the same classification procedure including only the ADHD GWAS. That is, we estimated the regression coefficient on a training dataset including 70% of data and tested on the subset that was left out. Our multivariate model did not outperform a univariate predictor. In fact, in the case of EA, the univariate predictor even performed slightly better (P= 0.004, Wilcoxon rank-sum test, Figure S5), indicating the PGS for additional traits introduce more noise than signal to the classifier.

Since the EA cohort drives the observed genetic association between PS and ADHD, we next explored the nature and robustness of the association in the EA cohort only. Comparing the extremes of the distribution, the top decile of PGS was associated with a nearly 2.5-fold increased risk for being assigned to the PS group compared to the lowest decile (OR=2.43 (1.71, 3.51) P=1.25e-06, Figure 2b). Globally, within EA the ADHD PGS explains about 1% of variation in case-control status, as measured by reduction in Nagelkerke R2. Moreover, the association is (i) robust across P-value thresholds (ii) not driven by subtle population stratification within the EA cohort, (ii) extends to the European-only version of the ADHD GWAS (see Supplementary Results for details).

### Developmental effects

We hypothesized that the association between PS and ADHD PGS would be strongest in the younger children and weaker closer to the typical age of onset of schizophrenia and other psychotic disorders. As predicted, we observe a significant interaction of ADHD PGS with age, with a stronger association for younger children (age 12 or younger), weakening in late adolescence (P=0.02 for the interaction term in the full model, Figure 2c). Given this decrease of association across age, we hypothesized an association with schizophrenia PGS in the older age group, but this was not the case (Supplementary Results).

We observed no interaction with sex (P=0.85): the association between males and females are near-identical, with the OR in males-only 1.23 (1.10,1.37) and females-only 1.25 (1.10,1.41).

### Phenotypic overlap

While evidence for shared genetic risk for (categorically defined) SCZ and ADHD is minimal (genetic correlation is r=0.11, se=0.04, P=0.001, LD-score regression(17)), ADHD and psychosis symptoms are known to co-occur in youth(5, 6). Indeed, across ancestries, we observe a strong phenotypic overlap between PS and ADHD symptoms in the PNC (see Supplementary Materials). In the total sample, 5% of youth satisfy the DSM criteria for lifetime ADHD (n=384; n=222 and n=114 within EA and AA, respectively). A large fraction of ADHD cases also endorse psychosis spectrum symptoms, and the majority of subjects in the PS group endorse ADHD symptoms (OR 2-4.7, Table 1).

**Table 1.** Univariate association of ADHD PGS with PS across ancestry groups. EA_noADHD and EA_noSymptoms denote the EA group after removing ADHD cases and subjects that endorse any of the symptoms in the ADHD screener, respectively.

| Group | N | %PS | OR | 2.5% | 97.5% | P |
| --- | --- | --- | --- | --- | --- | --- |
| All | 7008 | 19% | 1.12 | 1.05 | 1.18 | 0.0003 |
| EA | 4790 | 15% | 1.23 | 1.14 | 1.34 | 4.15*10<sup>-7</sup> |
| AA | 1746 | 29% | 0.98 | 0.88 | 1.06 | 0.65 |
| EA_noADHD | 4349 | 13% | 1.26 | 1.16 | 1.38 | 2.63*10<sup>-7</sup> |
| EA_noSymptoms | 2497 | 7% | 1.23 | 1.05 | 1.44 | 0.01 |

We additionally explored the overlap between ADHD and PS using a variety of phenotypic constructs, and observed strong phenotypic overlap across domains, and across ancestry groups (Supplementary Methods and Results). For example, we observe strong positive correlations between ADHD and PS domain scores (For all correlations, spearman rho is 0.25 < r <0.69 and P<10^−16^). Both higher inattention and hyperactivity scores are associated with an increased probability of being classified as PS, while at the same time higher PRIME and SOPS scores are associated with increased probability of meeting ADHD criteria, across ancestry groups (Supplementary Results and Figures S9-S12). The age interaction effect we observed at the genetic level is not consistently observed at the phenotypic level; e.g., the effect of answering “yes” to any of the ADHD screener questions on the probability of being classified as PS does not change during development (P=0.41 in EA, Supplementary results).

### Phenotypic overlap and genetic liability

Given the strong phenotypic overlap between PS and ADHD symptoms, we performed a series of follow-up analyses to determine the extent to which the observed genetic association is driven by this overlap. We tested the association after removing ADHD cases. We also removed the subset of youth that endorsed any ADHD symptoms from the screener. The latter analysis leaves only 2,497 EA subjects and reduces the percentage of PS cases from 15% to 7%, with a total n= 169 PS subjects. In both cases, despite the smaller sample sizes and the relative depletion of the number of cases, the effect of ADHD PGS on PS risk remains stable (Table 2, Supplementary results, Table S2).

**Table 2.** Clinical overlap between PS and ADHD across ancestry groups. Confusion matrices for overlap between PS and subjects meeting DSM criteria for ADHD (ADHD) and subjects that endorse any of the symptoms in the ADHD screener (ADHD_Symptoms), respectively. Colors indicate different ancestry groups All pink, EA blue, AA Orange.

|  |  | PS_ALL |  | PS_EA |  | PS_AA |  |
| --- | --- | --- | --- | --- | --- | --- | --- |
|  |  | 1 | 0 | 1 | 0 | 1 | 0 |
| ADHD | 1 | 151 | 233 | 83 | 139 | 50 | 64 |
|  | 0 | 1115 | 5220 | 584 | 3765 | 426 | 1080 |
| Overlap Statistic |  | OR 3.0 (2.4-3.7),<br>P<2.2*10<sup>-16</sup> |  | OR 3.8 (2.8-5.1)<br>P<2.2*10<sup>-16</sup> |  | OR 1.9 (1.3-2.9)<br>P= 0.0008 |  |
| ADHD_Symptoms | 1 | 1073 | 2515 | 536 | 1694 | 427 | 621 |
|  | 0 | 284 | 3120 | 169 | 2328 | 91 | 573 |
| Overlap Statistic |  | OR 4.7 ( 4.1-5.4),<br>P<2.2*10<sup>-16</sup> |  | OR 4.6 (3.6-5.3),<br>P<2.2*10<sup>-16</sup> |  | OR 4.3 (3.3-5.6),<br>P<2.2*10<sup>-16</sup> |  |

Thus, the association between increased ADHD liability and PS in youth does not appear to be driven by symptom overlap. Moreover, ADHD genetic risk is not only associated with PS as a categorical variable, but similarly with the psychosis severity scales, measured quantitatively (beta=0.47, P=0.0002 for PRIME and beta= 0.21, P= 1.68*10^−8^ for SOPS), and significantly increased in the subset of PS subjects that endorse hallucinations (n=258; OR=1.15 (1.02,1.30), P=0.02). As expected, the ADHD PGS is associated with ADHD status (OR 1.18, CI(1.04, 1.36), P=0.01). Conversely, and contrary to existing evidence (48), the schizophrenia PGS is not associated with ADHD status (P= 0.51).

Finally, we investigate the relationship of ADHD and psychosis phenotypes to substance use. In the PNC EA cohort, substance use information was available for 43% of the sample. In this sample, PS overlapped the use of alcohol, tobacco and marijuana only nominally (P>0001), and not cocaine or over the counter substance use (P>0.05). Including these as covariates in the model did not alter the estimated effect sizes (See Table S2).

### Replication in an independent cohort

We sought to replicate our finding of increased polygenic liability for ADHD in youth with psychotic spectrum symptoms in the North American Prodrome Longitudinal Study, Phase 2 (NAPLS2) cohort(49, 50). NAPLS2 is an eight-site longitudinal study of predictors and mechanisms of conversion to psychosis, and includes help-seeking adolescents and young adults at clinical high-risk (CHR) for psychosis (ages 12-35, with a median age of 18; N=328) as well as unaffected control subjects (N=216) (51).

As in our discovery sample, we observed increased ADHD risk in CHR youth compared to controls (OR=1.06 (1.01, 1.11) P= 0.02; in EA, OR= 1.09 (1.01, 1.18) P=0.03; N=124 and N=70 respectively). These effect sizes are similar to the ones observed in the >16 age group in the PNC EA cohort (Figure 2). However, there is no difference in ADHD PGS between CHR subjects who subsequently convert to psychosis versus those who do not (N=80 converters, N=248 non-converters; P=0.62).

## Discussion

Leveraging a large community-based sample, we sought to characterize genetic risk profiles for psychotic spectrum symptoms across childhood and adolescence. We applied recently developed multi-PGS prediction models as well univariate statistical tests, based on GWAS of multiple brain and behavioral traits. We observed a modest but robust association between broadly defined psychosis symptomatology and genetic liability for ADHD, but not for schizophrenia or any other psychiatric traits. This effect was only observed in participants of European ancestry, for whom those within the highest decile of ADHD genetic risk have an almost 2.5-fold increased likelihood of being in the PS group, compared to those with lowest ADHD polygenic scores. This association, while modest, holds even when excluding individuals who endorsed any ADHD symptoms, is strongest in children twelve years or younger, and diminishes closer to typical age of onset of schizophrenia. ADHD polygenic scores, were never before tested for association with symptoms of psychosis in youth. We therefore replicated our finding in an independent cohort of subjects at clinical high risk for psychosis.

Contrary to recent genetic evidence based on psychotic experiences in adulthood (21), psychosis spectrum symptoms in youth did not yield a general association with multiple psychiatric illnesses, but with ADHD specifically. While an association between schizophrenia genetic risk and psychosis symptoms has been reported in adults, the lack of such an association in our study is consistent with previous literature in adolescents (in a population sample of >5k genotyped youth aged 12-18, no such association was observed (22)).

In our study, the multivariate approach did not improve classification accuracy above a single trait association of ADHD. However, future efforts to improve risk scoring methodology and especially more powerful GWAS of related traits are likely to improve prediction as well.

As the discovery GWAS of ADHD included 55,374 children and adults (20,183 ADHD cases and 35,191 control subjects), ADHD genetic risk across all developmental ranges is captured in the downstream polygenic scores. An important avenue for future work will be to investigate if the association with general psychotic symptomatology is driven more by genetic liability present in children versus adults diagnosed with ADHD.

Despite similar phenotypic correlations across ancestry groups, the absence of any genetic association in youth of African ancestry highlights the need for increasing ethnic diversity in GWA studies. Because allele frequencies, linkage disequilibrium patterns, and effect sizes of common polymorphisms vary with ancestry, current common variant genetic findings do not translate well across populations (46, 47, 52). Our study offers further evidence that polygenic scores, at this point, have limited predictive power in non-European ethnic groups. As PGS scores are approaching clinical utility(53), the crucial equity issue that arises as a result from this discrepancy should not be taken lightly. Novel tools to generate risk scores across ancestries, such as by scoring only segments of the genome matching the GWAS population in admixed populations, may improve applicability of the risk scores across ancestries. Most importantly, however, larger samples from different ancestries are needed to begin to close this gap.

Based on a follow up study of PNC youth, about half of youth experiencing psychotic symptoms, symptoms persist or get worse over a two-year follow up, while even those whose psychotic symptoms remit exhibited comparatively higher symptom levels and lower functioning than typically developing youths(7). An important follow-up question is whether youth with psychotic symptoms that ultimately develop a psychotic disorder have different genetic characteristics than those who do not. Consistent with our present findings in the PNC, we recently found that subjects meeting psychosis risk syndrome criteria that do not develop psychosis in a two year follow-up in the NAPLS cohort do not have increased genetic risk for schizophrenia compared to controls. However, polygenic liability for schizophrenia is a predictor for conversion to overt psychosis(6). We now show, in the same NAPLS cohort, that ADHD PGS is also increased in CHR youth, but was not associated with conversion. While there are important clinical differences between PNC PS and NAPLS CHR cohorts (importantly, the latter consists of help-seeking youth, whereas PNC is a community sample), the consistency of our findings across cohorts confirms the robustness of the ADHD PGS association with psychosis symptoms.

Our findings shed light on the genetic architecture of psychosis symptoms in a population-based youth cohort and suggest that broad psychosis spectrum symptoms in youth may reflect a different genetic etiology than psychotic symptoms in adulthood, or those that convert to psychosis. Rather, the genetic etiology of psychosis symptoms in youth seems more akin to ADHD. Findings indicate that genetic risk can be investigated across early symptom trajectories, and in non-help seeking populations, to improve our understanding of disease risk factors and psychiatric comorbidities.

## Acknowledgements

This work was supported by R01MH107250 awarded to RAO and CB. LOL was supported by K99MH116115. Collaborative U01 award from the National Institute of Mental Health at the National Institutes of Health (MH081902 to TDC and CEB; MH081857 to BAC; MH081988 to EW; MH081928 to LS; MH082004 to DP; MH081944 to KC; MH081984 to JA; MH082022 to SWW; MH076989; MH107235 to RCG; MH081902 to REG). We thank all research participants and researchers involved in making each GWAS summary statistic available and this work possible, including the 23andMe Inc. Research Team.

## URLS

Ricopili https://sites.google.com/a/broadinstitute.org/ricopili/

## Supplementary Figure legends

**Figure 1.** A) Relative admixture ancestry components (based on K=3) for the EA and AA subjects in the PNC B) Cross-validation error for different ancestry components K.

**Figure 2.** MDS components based on a set of high quality independent SNPs. Orange indicates individuals assigned AA ancestry, light blue EA and dark blue are subjects with >95% EA, used for heritability analyses.

**Figure 3.** Identity by state estimates >0.05 between EA and AA pairs estimated from analysis including the whole cohort that were not identified in within-ancestry analyses. Nearly all pairs (∼98%) are pairs of AA subjects.

**Figure 4.** Mean regression coefficients for the multivariate PGS based on different GWAS in All, EA and AA subjects. Standard errors indicate standard errors of the mean. Actual data is displayed on the left, models based on case-control permutations on the right.

**Figure 5.** AUCROC for each elastic net model trained on 70% and tested on the remaining 30% of data in All (pink), EA (blue) and AA (orange). Box plots indicate the median and the lower and upper hinges correspond to the first and third quartiles. The grey dots and boxplots here refer to a model including only the ADHD PGS within each ancestry group.

**Figure 6.** Standardized ADHD PGS in youth with PS versus non-PS youth for All, EA, and AA.

**Figure 7.** Association of ADHD PGS with PS in EA varying the P-value threshold.

**Figure 8.** Correlation between ADHD PGS based on the entire sample and the European-only sample.

**Figure 9.** Spearman rank correlations between PRIME and SOPS psychosis scores and Inattention and hyperactivity ADHD scores in EA, AA, and All samples.

**Figure 10.** Relation between Inattention and Hyperactivity scores and proportion of PS cases in EA, AA, and All samples.

**Figure 11.** Relation between PRIME and SOPS scores and proportion of ADHD cases in EA, AA, and All samples.

**Figure 12.** Inattention (a), Hyperactivity (b), PRIME (c) and SOPS (d) scores by case category: neither (subjects that are not ADHD or PS cases), ADD_only (ADHD cases that are not PS cases), PS_only (PS cases that are not ADHD cases), both (subjects that are both PS and ADHD cases), in EA, AA, and All samples.

## Supplementary Table legends

**Table 1. Summary of GWAS used for analyses** For summary statistics not available through LD-hub, we estimated the heritability on the liability scale and other GWAS parameters by running LD score regression(17), using the population prevalence as reported in the GWAS.

**Table 2. Univariate association of ADHD PGS with PS across ancestry groups**, correcting for phenotypic overlap and substance use, where: 95 subjects with >95% EA; noADHD removing ADHD subjects; noADD011 removing subjects that answer “yes” to the question ADD011 “Did you often have trouble paying attention or keeping your mind on your school, work, chores, or other activities that you were doing?”; Hyperactivityscore is the Hyperactivity score, InattentionScore is the Inattention score; noSymptoms removing subjects that endorse any of the symptoms in the ADHD screener; Substances: OTC over the counter, ALC alcohol, TOB tobacco, MAR marihuana, COC cocaine.

## References

1. McGrath JJ, Saha S, Al-Hamzawi A, Alonso J, Bromet EJ, Bruffaerts R, et al. Psychotic Experiences in the General Population: A Cross-National Analysis Based on 31,261 Respondents From 18 Countries. JAMA Psychiatry. 2015;72(7):697–705.

2. Saha S, Chant D, Welham J, McGrath J. A systematic review of the prevalence of schizophrenia. PLoS Med. 2005;2(5):e141.

3. Kessler RC, Chiu WT, Demler O, Merikangas KR, Walters EE. Prevalence, severity, and comorbidity of 12-month DSM-IV disorders in the National Comorbidity Survey Replication. Arch Gen Psychiatry. 2005;62(6):617–27.

4. Schimmelmann BG, Michel C, Martz-Irngartinger A, Linder C, Schultze-Lutter F. Age matters in the prevalence and clinical significance of ultra-high-risk for psychosis symptoms and criteria in the general population: Findings from the BEAR and BEARS-kid studies. World Psychiatry. 2015;14(2):189–97.

5. Lee TY, Lee J, Kim M, Choe E, Kwon JS. Can We Predict Psychosis Outside the Clinical High-Risk State? A Systematic Review of Non-Psychotic Risk Syndromes for Mental Disorders. Schizophr Bull. 2018;44(2):276–85.

6. Addington J, Piskulic D, Liu L, Lockwood J, Cadenhead KS, Cannon TD, et al. Comorbid diagnoses for youth at clinical high risk of psychosis. Schizophr Res. 2017;190:90–5.

7. Calkins ME, Moore TM, Satterthwaite TD, Wolf DH, Turetsky BI, Roalf DR, et al. Persistence of psychosis spectrum symptoms in the Philadelphia Neurodevelopmental Cohort: a prospective two-year follow-up. World Psychiatry. 2017;16(1):62–76.

8. Caspi A, Moffitt TE. All for One and One for All: Mental Disorders in One Dimension. Am J Psychiatry. 2018;175(9):831–44.

9. Hartmann JA, Nelson B, Ratheesh A, Treen D, McGorry PD. At-risk studies and clinical antecedents of psychosis, bipolar disorder and depression: a scoping review in the context of clinical staging. Psychol Med. 2019;49(2):177–89.

10. Bufferd SJ, Dougherty LR, Carlson GA, Rose S, Klein DN. Psychiatric disorders in preschoolers: continuity from ages 3 to 6. Am J Psychiatry. 2012;169(11):1157–64.

11. Fisher HL, Caspi A, Poulton R, Meier MH, Houts R, Harrington H, et al. Specificity of childhood psychotic symptoms for predicting schizophrenia by 38 years of age: a birth cohort study. Psychol Med. 2013;43(10):2077–86.

12. Schizophrenia Working Group of the Psychiatric Genomics C. Biological insights from 108 schizophrenia-associated genetic loci. Nature. 2014;511(7510):421–7.

13. Stahl EA, Breen G, Forstner AJ, McQuillin A, Ripke S, Trubetskoy V, et al. Genome-wide association study identifies 30 loci associated with bipolar disorder. Nat Genet. 2019;51(5):793–803.

14. Maier RM, Visscher PM, Robinson MR, Wray NR. Embracing polygenicity: a review of methods and tools for psychiatric genetics research. Psychol Med. 2018;48(7):1055–67.

15. Bergen SE, Ploner A, Howrigan D, Group CNVA, the Schizophrenia Working Group of the Psychiatric Genomics C, O’Donovan MC, et al. Joint Contributions of Rare Copy Number Variants and Common SNPs to Risk for Schizophrenia. Am J Psychiatry. 2019;176(1):29–35.

16. Allardyce J, Leonenko G, Hamshere M, Pardinas AF, Forty L, Knott S, et al. Association Between Schizophrenia-Related Polygenic Liability and the Occurrence and Level of Mood-Incongruent Psychotic Symptoms in Bipolar Disorder. JAMA Psychiatry. 2018;75(1):28–35.

17. Bulik-Sullivan BK, Loh PR, Finucane HK, Ripke S, Yang J, Schizophrenia Working Group of the Psychiatric Genomics C, et al. LD Score regression distinguishes confounding from polygenicity in genome-wide association studies. Nat Genet. 2015;47(3):291–5.

18. Pain O, Dudbridge F, Cardno AG, Freeman D, Lu Y, Lundstrom S, et al. Genome-wide analysis of adolescent psychotic-like experiences shows genetic overlap with psychiatric disorders. Am J Med Genet B Neuropsychiatr Genet. 2018;177(4):416–25.

19. Zavos HM, Freeman D, Haworth CM, McGuire P, Plomin R, Cardno AG, et al. Consistent etiology of severe, frequent psychotic experiences and milder, less frequent manifestations: a twin study of specific psychotic experiences in adolescence. JAMA Psychiatry. 2014;71(9):1049–57.

20. Sieradzka D, Power RA, Freeman D, Cardno AG, Dudbridge F, Ronald A. Heritability of Individual Psychotic Experiences Captured by Common Genetic Variants in a Community Sample of Adolescents. Behav Genet. 2015;45(5):493–502.

21. Legge SE, Jones HJ, Kendall KM, Pardiñas AF, Menzies G, Bracher-Smith M, et al. Genetic association study of psychotic experiences in UK Biobank. BioRxiv. 2019:583468.

22. Jones HJ, Stergiakouli E, Tansey KE, Hubbard L, Heron J, Cannon M, et al. Phenotypic Manifestation of Genetic Risk for Schizophrenia During Adolescence in the General Population. JAMA Psychiatry. 2016;73(3):221–8.

23. Satterthwaite TD, Vandekar SN, Wolf DH, Bassett DS, Ruparel K, Shehzad Z, et al. Connectome-wide network analysis of youth with Psychosis-Spectrum symptoms. Mol Psychiatry. 2015;20(12):1508–15.

24. Satterthwaite TD, Wolf DH, Calkins ME, Vandekar SN, Erus G, Ruparel K, et al. Structural Brain Abnormalities in Youth With Psychosis Spectrum Symptoms. JAMA Psychiatry. 2016;73(5):515–24.

25. Jalbrzikowski M, Freedman D, Hegarty CE, Mennigen E, Karlsgodt KH, Olde Loohuis LM, et al. Structural Brain Alterations in Youth With Psychosis and Bipolar Spectrum Symptoms. J Am Acad Child Adolesc Psychiatry. 2019.

26. Gur RC, Calkins ME, Satterthwaite TD, Ruparel K, Bilker WB, Moore TM, et al. Neurocognitive growth charting in psychosis spectrum youths. JAMA Psychiatry. 2014;71(4):366–74.

27. Krapohl E, Patel H, Newhouse S, Curtis CJ, von Stumm S, Dale PS, et al. Multi-polygenic score approach to trait prediction. Mol Psychiatry. 2018;23(5):1368–74.

28. Satterthwaite TD, Connolly JJ, Ruparel K, Calkins ME, Jackson C, Elliott MA, et al. The Philadelphia Neurodevelopmental Cohort: A publicly available resource for the study of normal and abnormal brain development in youth. Neuroimage. 2016;124(Pt B):1115–9.

29. Kobayashi H, Nemoto T, Koshikawa H, Osono Y, Yamazawa R, Murakami M, et al. A self-reported instrument for prodromal symptoms of psychosis: testing the clinical validity of the PRIME Screen-Revised (PS-R) in a Japanese population. Schizophr Res. 2008;106(2-3):356–62.

30. Kaufman J, Birmaher B, Brent D, Rao U, Flynn C, Moreci P, et al. Schedule for Affective Disorders and Schizophrenia for School-Age Children-Present and Lifetime Version (K-SADS-PL): initial reliability and validity data. J Am Acad Child Adolesc Psychiatry. 1997;36(7):980–8.

31. Miller TJ, McGlashan TH, Rosen JL, Cadenhead K, Cannon T, Ventura J, et al. Prodromal assessment with the structured interview for prodromal syndromes and the scale of prodromal symptoms: predictive validity, interrater reliability, and training to reliability. Schizophr Bull. 2003;29(4):703–15.

32. Calkins ME, Moore TM, Merikangas KR, Burstein M, Satterthwaite TD, Bilker WB, et al. The psychosis spectrum in a young U.S. community sample: findings from the Philadelphia Neurodevelopmental Cohort. World Psychiatry. 2014;13(3):296–305.

33. Alexander DH, Novembre J, Lange K. Fast model-based estimation of ancestry in unrelated individuals. Genome Res. 2009;19(9):1655–64.

34. Zheng J, Erzurumluoglu AM, Elsworth BL, Kemp JP, Howe L, Haycock PC, et al. LD Hub: a centralized database and web interface to perform LD score regression that maximizes the potential of summary level GWAS data for SNP heritability and genetic correlation analysis. Bioinformatics. 2017;33(2):272–9.

35. Demontis D, Walters RK, Martin J, Mattheisen M, Als TD, Agerbo E, et al. Discovery of the first genome-wide significant risk loci for attention deficit/hyperactivity disorder. Nat Genet. 2019;51(1):63–75.

36. Autism Spectrum Disorders Working Group of The Psychiatric Genomics C. Meta-analysis of GWAS of over 16,000 individuals with autism spectrum disorder highlights a novel locus at 10q24.32 and a significant overlap with schizophrenia. Mol Autism. 2017;8:21.

37. Cross-Disorder Group of the Psychiatric Genomics C. Identification of risk loci with shared effects on five major psychiatric disorders: a genome-wide analysis. Lancet. 2013;381(9875):1371–9.

38. Wray NR, Ripke S, Mattheisen M, Trzaskowski M, Byrne EM, Abdellaoui A, et al. Genome-wide association analyses identify 44 risk variants and refine the genetic architecture of major depression. Nat Genet. 2018;50(5):668–81.

39. Hyde CL, Nagle MW, Tian C, Chen X, Paciga SA, Wendland JR, et al. Identification of 15 genetic loci associated with risk of major depression in individuals of European descent. Nat Genet. 2016;48(9):1031–6.

40. consortium C. Sparse whole-genome sequencing identifies two loci for major depressive disorder. Nature. 2015;523(7562):588–91.

41. Hibar DP, Stein JL, Renteria ME, Arias-Vasquez A, Desrivieres S, Jahanshad N, et al. Common genetic variants influence human subcortical brain structures. Nature. 2015;520(7546):224–9.

42. Hu Y, Shmygelska A, Tran D, Eriksson N, Tung JY, Hinds DA. GWAS of 89,283 individuals identifies genetic variants associated with self-reporting of being a morning person. Nat Commun. 2016;7:10448.

43. Okbay A, Baselmans BM, De Neve JE, Turley P, Nivard MG, Fontana MA, et al. Corrigendum: Genetic variants associated with subjective well-being, depressive symptoms, and neuroticism identified through genome-wide analyses. Nat Genet. 2016;48(12):1591.

44. Vilhjalmsson BJ, Yang J, Finucane HK, Gusev A, Lindstrom S, Ripke S, et al. Modeling Linkage Disequilibrium Increases Accuracy of Polygenic Risk Scores. Am J Hum Genet. 2015;97(4):576–92.

45. Zou H, Hastie T. Regularization and variable selection via the elastic net. Journal of the royal statistical society: series B (statistical methodology). 2005;67(2):301–20.

46. Mills MC, Rahal C. A scientometric review of genome-wide association studies. Commun Biol. 2019;2:9.

47. Martin AR, Gignoux CR, Walters RK, Wojcik GL, Neale BM, Gravel S, et al. Human Demographic History Impacts Genetic Risk Prediction across Diverse Populations. Am J Hum Genet. 2017;100(4):635–49.

48. Nivard MG, Gage SH, Hottenga JJ, van Beijsterveldt CEM, Abdellaoui A, Bartels M, et al. Genetic Overlap Between Schizophrenia and Developmental Psychopathology: Longitudinal and Multivariate Polygenic Risk Prediction of Common Psychiatric Traits During Development. Schizophr Bull. 2017;43(6):1197–207.

49. Cannon TD, Yu C, Addington J, Bearden CE, Cadenhead KS, Cornblatt BA, et al. An Individualized Risk Calculator for Research in Prodromal Psychosis. Am J Psychiatry. 2016;173(10):980–8.

50. Addington J, Liu L, Buchy L, Cadenhead KS, Cannon TD, Cornblatt BA, et al. North American Prodrome Longitudinal Study (NAPLS 2): The Prodromal Symptoms. J Nerv Ment Dis. 2015;203(5):328–35.

51. Diana O. Perkins LOL, Jenna Barbee, John Ford, Clark D. Jeffries, Jean Addington, Carrie E. Bearden, Kristin S. Cadenhead, Tyrone D. Cannon, Barbara A. Cornblatt, Daniel H. Mathalon, Daniel H. Mathalon, Larry J. Seidman, Ming Tsuang, Ming Tsuang Polygenic Risk Score Contribution to Psychosis Prediction in a Target Population of Persons at Clinical High-Risk. American Journal of Psychiatry 2019 To Appear.

52. Martin AR, Kanai M, Kamatani Y, Okada Y, Neale BM, Daly MJ. Clinical use of current polygenic risk scores may exacerbate health disparities. Nat Genet. 2019;51(4):584–91.

53. Khera AV, Chaffin M, Aragam KG, Haas ME, Roselli C, Choi SH, et al. Genome-wide polygenic scores for common diseases identify individuals with risk equivalent to monogenic mutations. Nat Genet. 2018;50(9):1219–24.

