## Supplementary Note for "Genetic and clinical analyses of psychosis spectrum symptoms in a large multi-ethnic youth cohort reveal significant link with ADHD"

**Supplementary Methods**

Definition of psychosis spectrum

Criteria to establish a group of individuals that experiences psychosis spectrum symptoms were defined as in prior PNC studies(1-4). Specifically, positive psychotic symptoms were assessed with items from the PRIME Screen Revised (PRIME S-R) as well as the KSADS and negative/disorganized symptoms were assessed with items from the Scale of Prodromal Symptoms (SOPS).

PS participants fell into at least one of the following categories:

- PRIME Screen-Revised (PS-R) based criteria assessing positive symptoms:
  a. one item rated as 6 (“definitely agree”) OR
  b. three or more items rated as 5 (“somewhat agree”) OR
  c. the total score of all PS-R items was above 2 standard deviations (SD) of the participant’s age group’s mean. The rationale for this is to determine whether symptoms endorsed would be considered developmentally inappropriate, given the subject’s age. Previous publications have described the clinical and functional significance of this cut-off.
- K-SADS criterion assessing lifetime hallucinations and delusional symptoms:
  a. endorsement of definite or possible hallucinations or delusions lasting at least one day outside the context of substance use and physical illness but accompanied by significant distress (rating of >= 5).
- Scale of prodromal symptoms (SOPS) criterion assessing negative and disorganized symptoms:

the total score of SOPS negative and disorganized items was 2 SDs above the mean for the participant’s age group. The rationale for this is to determine whether symptoms endorsed would be considered developmentally inappropriate, given the subject’s age.

Description of ADHD symptoms.

Criteria that we applied to establish ADHD categorical diagnosis were based on a previous study on the PNC data in order to provide consistency(5). Specifically, participants had to endorse inattentive or hyperactive symptoms that occurred in two or more settings, bothered the participant significantly, and that were present at the time of the assessment. In addition to a single unified category of ADHD, we included two rating scores as well as a binary phenotypes resulting from the following questions: 'ADD011' 'ADD012' 'ADD013' 'ADD014' 'ADD015' 'ADD016' 'ADD017' 'ADD018' 'ADD019' 'ADD020' 'ADD021' 'ADD022' 'ADD023' 'ADD024' 'ADD025' 'ADD032' 'ADD034a' 'ADD035' 'ADD050'. The sum of inattention symptoms and hyperactivity symptoms were used in follow-up phenotypic analyses. Since these numeric scores have few values (4 and 7 respectively) correlations with other trait variables were compared using spearman rank correlations. The question ADD011 is: “Did you often have trouble paying attention or keeping your mind on your school, work, chores, or other activities that you were doing?”

Genotyping QC and imputation

Genotyping was performed on multiple arrays: The vast majority of the samples were genotyped on the 550HH and 610Q SNP arrays from Illumina.  Quality control and imputation was performed using 1KG by the Broad Institute in two batches (n=1,652 and n=6,122). Imputation followed the standard Ricopili pipeline (see urls) and best-guess genotypes of well-imputed variants (INFO>0.8) were selected for further analysis. After merging the data from different imputation batches, we filtered on a genotyping rate >98%, minor allele frequency (MAF) of > 1%, resulting in a total of 1,937,561 SNPS for 7,774 individuals, 7,764 with phenotyping data.

Selection of EA and AA individuals

To identify individuals of European and African ancestry genetically, we used ADMIXTURE(1), a tool developed to estimate genetic ancestry from genotyping data. Based on cross-validation, we selected a k=3 as the number of global ancestral populations (see Supplementary Figure 1). We identified individuals of European ancestry (EA) based on Anc2>0.7 and African Ancestry (AA) based on Anc3>0.7. These settings resulted in a high concordance with self-reported ethnicity: 99% of individuals identified as European ancestry were included in the EA cohort, compared to 91% for AA. Principal components were computed based on a set of high quality independent SNPS (plink settings --indep-pairwise 50 5 0.2 --hwe 0.001 --mind 0.05), excluding highly variable regions on chromosomes 5,6,8, and 11 (n=115,869 SNPs). These principal components cluster EA and AA samples as expected (Figure 1, Figure S2). A total of 5,162 and 2,002 individuals were selected as EA, and AA respectively, with the remaining 600 included in neither category.

Removal of related individuals

We tested the sample for relatedness of individuals. The PNC cohort includes 477 sibling pairs related at 0.4 < pi-hat <0.6.  We observed that identity by state estimates may be influenced by LD clumping, which can inflate relatedness estimates, especially in a multi-ethnic cohort. We therefore computed pi-hat within EA and AA ancestries separately, removing at random one individual of each related pair related at a level >0.2 (level of cousin).  For those pairs discordant for ancestry (EA, AA, other) we determined relatedness based pi-hat estimates from the entire cohort (also using a cutoff of >0.2). In our sample of EA and AA individuals, we observed many identity by state relations in the global analyses compared to the within-ancestry, nearly always involving AA pairs, the minority ancestry in our sample (n>5,000 at pi-hat >0.05, with >98% AA-AA pairs). However, in our sample, these relationships reach at most a PI_HAT of 0.07 (See Figure S3). Since relatedness criteria are often used as a QC step to identify sample contamination, we caution that within-sample ancestry differences need to be taken into account in these analyses. After these final filtering steps, the sample sizes were All = 7,225, EA=4,852 and AA=1,802.

Heritability analyses

Heritability of psychosis spectrum symptoms was estimated using GCTA’s REML(6). Because heritability estimates are especially sensitive to population structure and cannot be estimated in a multi-ethnic cohort, we estimated heritability in only those unrelated individuals with >95% EA ancestry as measure from Admixture (n=4,138, with 593 cases and 3545 controls, Figure S3). Prior to performing GCTA, we removed SNPs in high LD (using --indep-pairwise 100kb 5 0.3), resulting in 125,318 independent SNPs. We included the first 10 genotyping PCs, imputation batch, sex, age and age^2^ into our heritability analyses. Heritability estimates are reported on the liability scale with a prevalence of 15%.

Polygenic scoring

To perform polygenic scoring we performed QC on the GWAS summary statistic files using the munge_sumstats.py pipeline from LDscore regression which filters on INFO > 0.9, MAF > 0.01 and 0 < P <= 1(7). We also removed variants that are not SNPs (e.g., indels), strand ambiguous SNPs, and duplicated SNPs. Clumping was performed based on 500Kb and r2=0.25 using the PNC dataset as a reference. For ancestry specific analyses, clumping was performed within each sub-cohort.  Polygenic scoring was performed using PLINK18 based on a p-value threshold of 0.05 for each GWAS. Depending on the trait, we used either the regression coefficient beta (for quantitative traits) of the log odds ratio (binary traits) as weights in the scoring algorithm. Prior to follow up analyses we corrected PRS scores for 10 principal components as well as imputation batch, and scaled the residuals to z-scores. For global analyses involving the entire sample we included PCs computed based on the full dataset.  In case of analyses only involving the EA (or AA) cohort, we included EA (or AA) specific PCs after exclusion of related samples. The standardized residuals were used for follow-up analyses. For follow up analyses we computed PRS for a range of p-value thresholds (5x10^-8^, 1x10^06^, 1x10^-4^, 0.001, 0.01, 0.05, 0.1, 0.2, 0.5, 1.0).

Replication in the NAPLS2 cohorts

The North American Prodrome Longitudinal Study, Phase 2 (NAPLS2)(8-10) is a two-year, eight-site study of predictors and mechanisms of conversion to psychosis. The study included 763 high-risk and 279 unaffected subjects, mostly adolescents and young adults (mean age 12-35 with a median age of 18). After removing high-risk subjects that did not meet COPS criteria, subjects without DNA samples, high-risk non-converters who did not complete the 2-year study, and 1 randomly selected sibling of 17 sibling pairs, a total of 328 clinical high risk subjects and 216 unaffected subjects with genetic data were included after QC (see (11) for details). Of the 328 clinical high risk subjects, a total of 80 converted to psychosis during the study. ADHD PRS was computed as described above.

**Supplementary Results**

**Ancestry**

Since polygenic risk analyses have been shown to be sensitive to slight population stratification, we added the Ancestry components to the regression, which did not have an effect (P> 0.4) or alter the association (OR 1.23 (1.137,1.34), P=4.40e-07). No association was observed between Anc2 and PS in logistic regression (P=0.61), nor with the ADHD PRS (P=0.95). We also checked whether Anc2 (the European ancestry component) interacts with age in predicting PS, but it did not (P= 0.62). Anc2 alone was only very modestly associated with age in a linear regression model (P=0.03). Finally, we tested the association in the group with >95% European ancestry, which again showed a similar effect (OR 1.25 (1.14, 1.37) P= 8.67e-07)

In addition to the main GWAS summary statistics, the ADHD PGC also provides a version including only samples of European ancestry, consisting of largely the same sample(12). The PRS based on the two summary statistics were near-identical, as well as the relative association (Figure S8).

**Heritability**

In addition to estimating confidence intervals using GCTA’s REML, we estimated confidence intervals of the heritability using FIESTA(13), which did not yield more precise results.

**ADHD two different summary statistics**

We observed a strong correlation between the two PRS generated from the full ADHD GWAS and the European only subset: R=0.92, P< 2.2e-16 (P=0.05).

**Schizophrenia PRS association by age**

We hypothesized that there may be an association with schizophrenia liability in the older age group (>16), but this was not the case (P= 0.31).

**Phenotypic overlap**

To explore the phenotypic overlap between ADHD and PS we assessed the correlation (measured using Spearman rank correlation) between inattention and hyperactivity scores for ADHD, and PRIME and SOPS scores for PS across ancestries (Figure S9, all correlations have P<10^-16^). Both higher inattention and hyperactivity scores are associated with increased proportion of PS cases(Figure S10)., while at the same time PRIME and SOPS scores are strongly correlated with higher proportions of ADHD (Figure S11). However, directly comparing these four dimensional scores across groups, we observe that ADHD and PS are distinct groups behaving similarly across ancestries (Figure S12). We checked for an interaction with age and the association between ADHD phenotypes as PS, and observe no consistent effect: the effect of answering “yes” to any of the ADHD screener questions does not interact significantly with age (P=0.41 in EA); ADD011 nomically interacted with age (P=0.02, with the association decreasing with increased age); neither association of PS with inattention of hyperactivity scores interacted with age (P=0.63, P=0.45 respectively). In AA, no interaction was observed.

**Phenotypic overlap and genetic risk**

We tested the associated after: removing ADHD cases, removing all youth that do not endorse a single ADHD symptom from the screener, correcting for inattention and hyperactivity scores. All results show robust effect size (Supplementary Table 2).

1.	Alexander DH, Novembre J, Lange K. Fast model-based estimation of ancestry in unrelated individuals. Genome Res. 2009;19(9):1655-64.

2.	Satterthwaite TD, Wolf DH, Calkins ME, Vandekar SN, Erus G, Ruparel K, et al. Structural Brain Abnormalities in Youth With Psychosis Spectrum Symptoms. JAMA Psychiatry. 2016;73(5):515-24.

3.	Calkins ME, Moore TM, Satterthwaite TD, Wolf DH, Turetsky BI, Roalf DR, et al. Persistence of psychosis spectrum symptoms in the Philadelphia Neurodevelopmental Cohort: a prospective two-year follow-up. World Psychiatry. 2017;16(1):62-76.

4.	Calkins ME, Moore TM, Merikangas KR, Burstein M, Satterthwaite TD, Bilker WB, et al. The psychosis spectrum in a young U.S. community sample: findings from the Philadelphia Neurodevelopmental Cohort. World Psychiatry. 2014;13(3):296-305.

5.	Kaufman J, Birmaher B, Brent D, Rao U, Flynn C, Moreci P, et al. Schedule for Affective Disorders and Schizophrenia for School-Age Children-Present and Lifetime Version (K-SADS-PL): initial reliability and validity data. J Am Acad Child Adolesc Psychiatry. 1997;36(7):980-8.

6.	Yang J, Lee SH, Goddard ME, Visscher PM. GCTA: a tool for genome-wide complex trait analysis. Am J Hum Genet. 2011;88(1):76-82.

7.	Bulik-Sullivan BK, Loh PR, Finucane HK, Ripke S, Yang J, Schizophrenia Working Group of the Psychiatric Genomics C, et al. LD Score regression distinguishes confounding from polygenicity in genome-wide association studies. Nat Genet. 2015;47(3):291-5.

8.	Hibar DP, Stein JL, Renteria ME, Arias-Vasquez A, Desrivieres S, Jahanshad N, et al. Common genetic variants influence human subcortical brain structures. Nature. 2015;520(7546):224-9.

9.	Cannon TD, Yu C, Addington J, Bearden CE, Cadenhead KS, Cornblatt BA, et al. An Individualized Risk Calculator for Research in Prodromal Psychosis. Am J Psychiatry. 2016;173(10):980-8.

10.	Addington J, Liu L, Buchy L, Cadenhead KS, Cannon TD, Cornblatt BA, et al. North American Prodrome Longitudinal Study (NAPLS 2): The Prodromal Symptoms. J Nerv Ment Dis. 2015;203(5):328-35.

11.	Diana O. Perkins LOL, Jenna Barbee, John Ford, Clark D. Jeffries, Jean Addington , Carrie E. Bearden, Kristin S. Cadenhead, Tyrone D. Cannon, Barbara A. Cornblatt, Daniel H. Mathalon, Daniel H. Mathalon, Larry J. Seidman, Ming Tsuang, Ming Tsuang Polygenic Risk Score Contribution to Psychosis Prediction in a Target Population of Persons at Clinical High-Risk. American Journal of Psychiatry 2019 To Appear.

12.	Demontis D, Walters RK, Martin J, Mattheisen M, Als TD, Agerbo E, et al. Discovery of the first genome-wide significant risk loci for attention deficit/hyperactivity disorder. Nat Genet. 2019;51(1):63-75.

13.	Schweiger R, Fisher E, Rahmani E, Shenhav L, Rosset S, Halperin E. Using stochastic approximation techniques to efficiently construct confidence intervals for heritability. Journal of Computational Biology. 2018;25(7):794-808.
