## Supplementary material for "Genetic and clinical analyses of psychosis spectrum symptoms in a large multi-ethnic youth cohort reveal significant link with ADHD": Table S2

Supplementary Table 2.

|  | **Group** | **N** | **%PS** | **OR** | **2.5%** | **97.5%** | **P** |
| --- | --- | --- | --- | --- | --- | --- | --- |
| **Ancestry analyses** | All | 7008 | 19% | 1.12 | 1.05 | 1.18 | 0.0003 |
|  | EA | 4790 | 15% | 1.23 | 1.14 | 1.34 | 4.15*10^-7^ |
|  | AA | 1746 | 29% | 0.98 | 0.88 | 1.06 | 0.65 |
|  | EA_95 | 4089 | 14% | 1.25 | 1.14 | 1.37 | 8.67*10^-7^ |
| **ADHD phenotypic overlap** | EA_noADHD | 4349 | 13% | 1.26 | 1.16 | 1.38 | 2.63*10^-7^ |
|  | EA_noADD011 | 2982 | 9% | 1.19 | 1.04 | 1.35 | 0.009 |
|  | EA_corrHyperactibityscore | 4647 | 15% | 1.19 | 1.09 | 1.29 | 8.25*10^-5^ |
|  | EA_corrInattentionscore | 4684 | 15% | 1.16 | 1.06 | 1.26 | 0.0007 |
|  | EA_noSymptoms | 2497 | 7% | 1.23 | 1.05 | 1.44 | 0.01 |
| **Substance use** | EA_corrSUBS_OTC | 2129 | 13% | 1.18 | 1.05 | 1.33 | 0.0058 |
|  | EA_corrSUBS_ALC | 2129 | 13% | 1.18 | 1.05 | 1.33 | 0.0066 |
|  | EA_corrSUBS_TOB | 2129 | 13% | 1.18 | 1.05 | 1.33 | 0.0070 |
|  | EA_corrSUBS_MAR | 2129 | 13% | 1.21 | 1.06 | 1.37 | 0.0061 |
|  | EA_corrSUBS_COC | 2129 | 13% | 1.19 | 1.05 | 1.34 | 0.0052 |
