## Supplementary Figures for "Genetic and clinical analyses of psychosis spectrum symptoms in a large multi-ethnic youth cohort reveal significant link with ADHD"

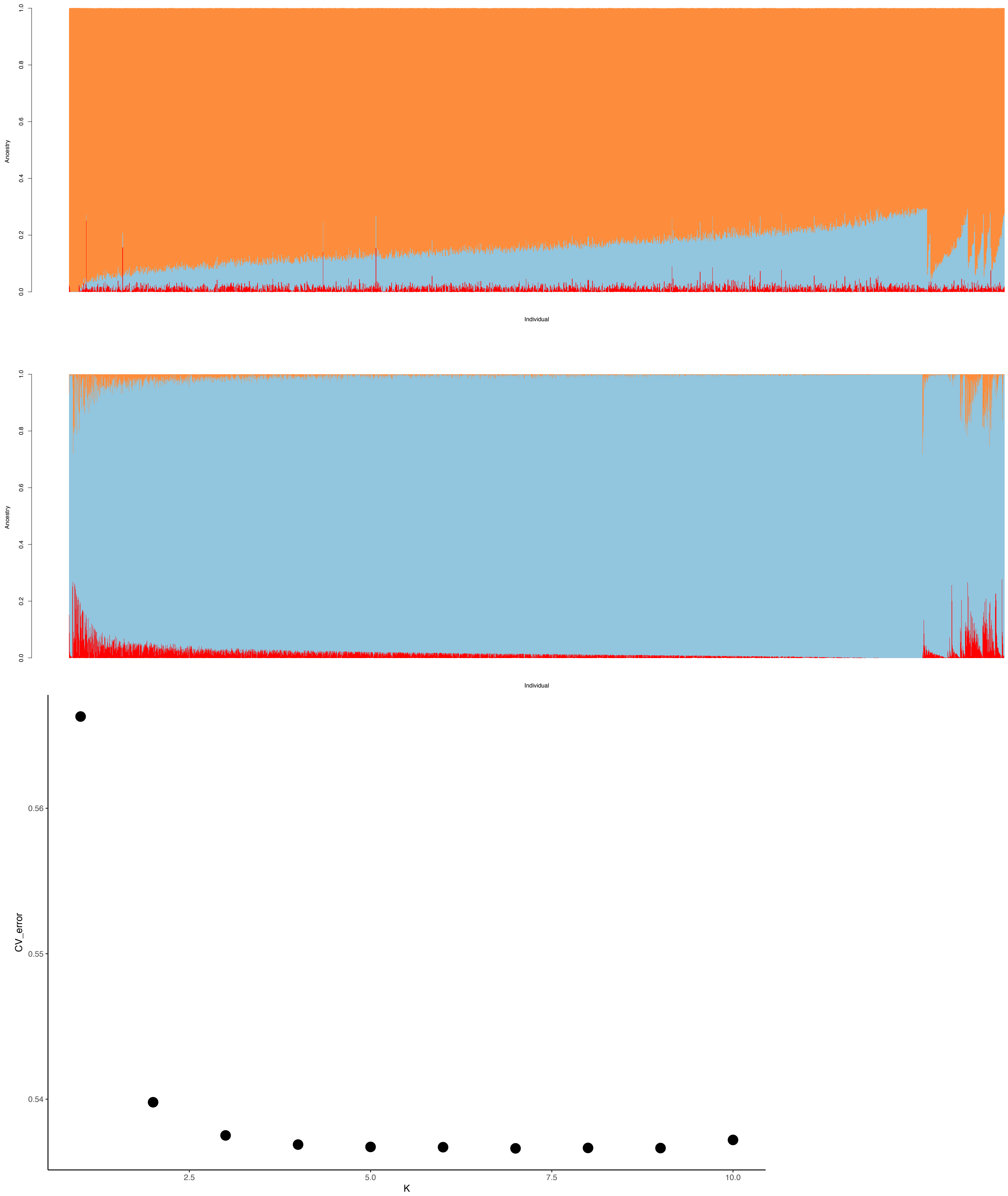
Ancestry
0.0
0.2
0.4
0.6
0.8
1.0
Individual
Ancestry
0.0
0.2
0.4
0.6
0.8
1.0
Individual
CV_error
0.54
0.55
0.56
2.5
5.0
7.5
10.0
K

**S Fig 1**

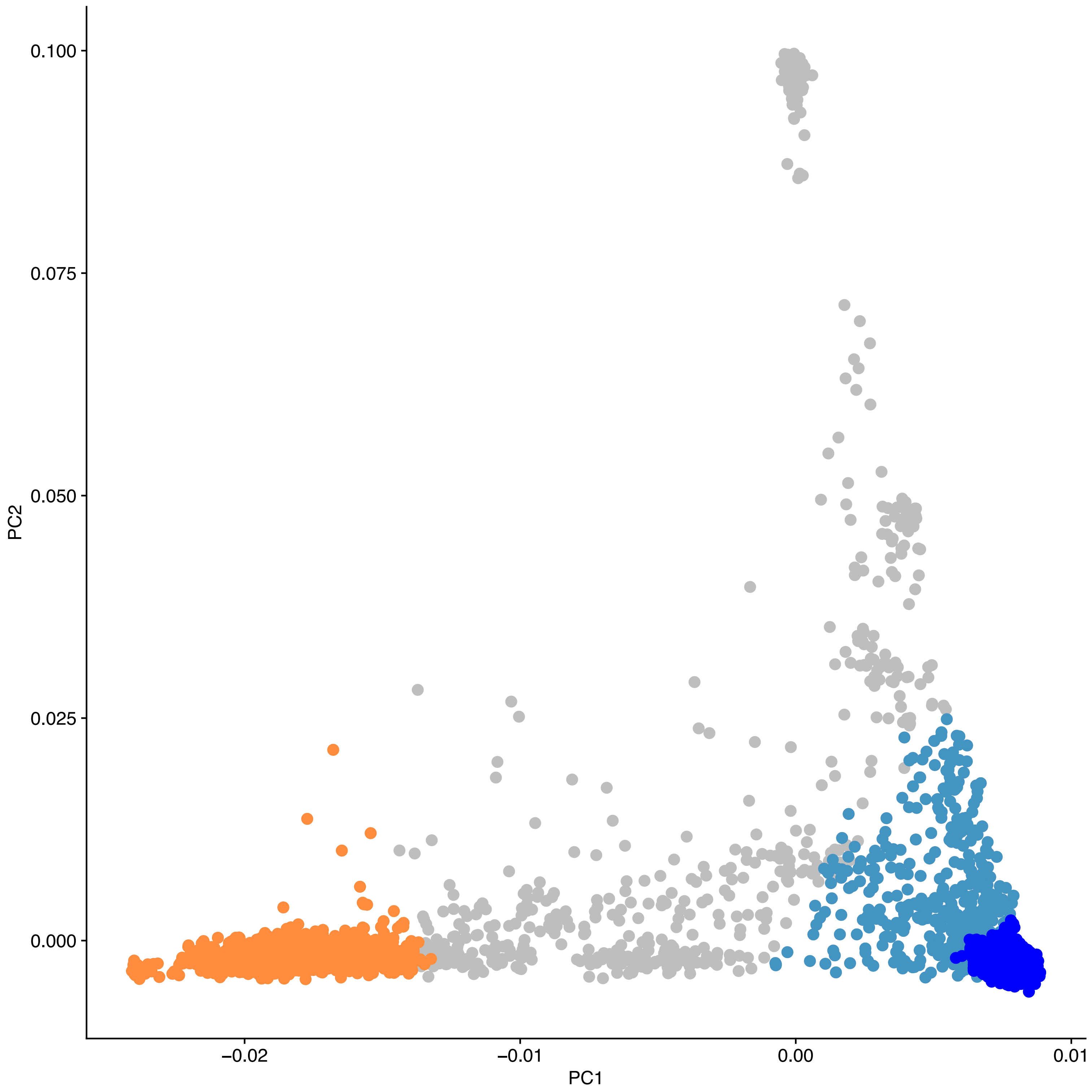


**S Fig 2**

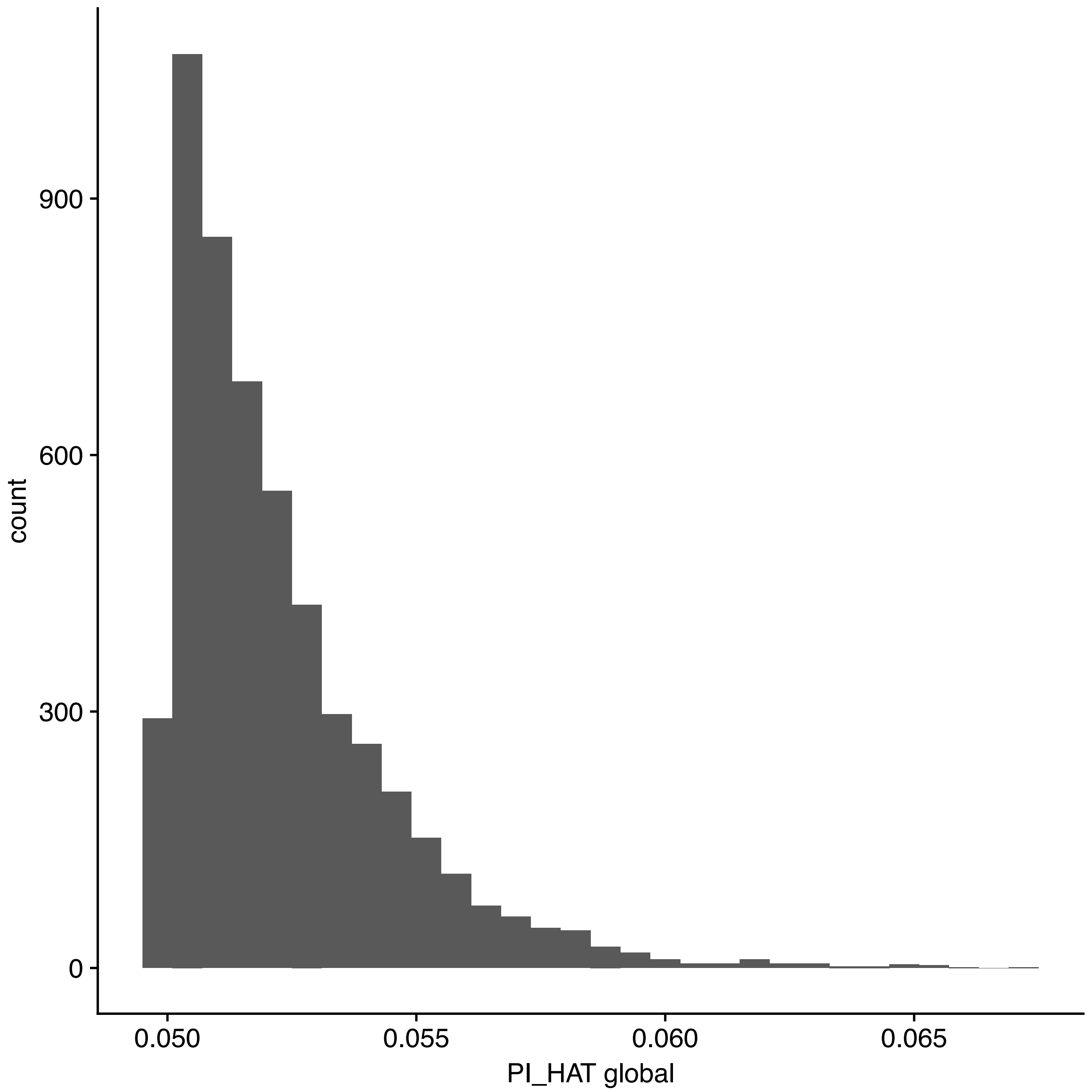


Identity by state estimates >0.05 between EA and AA pairs estimated from a global analyses that were not identified in within-ancestry analyses. Nearly all pairs (~98%) were AA pairs.

**S Fig 3**

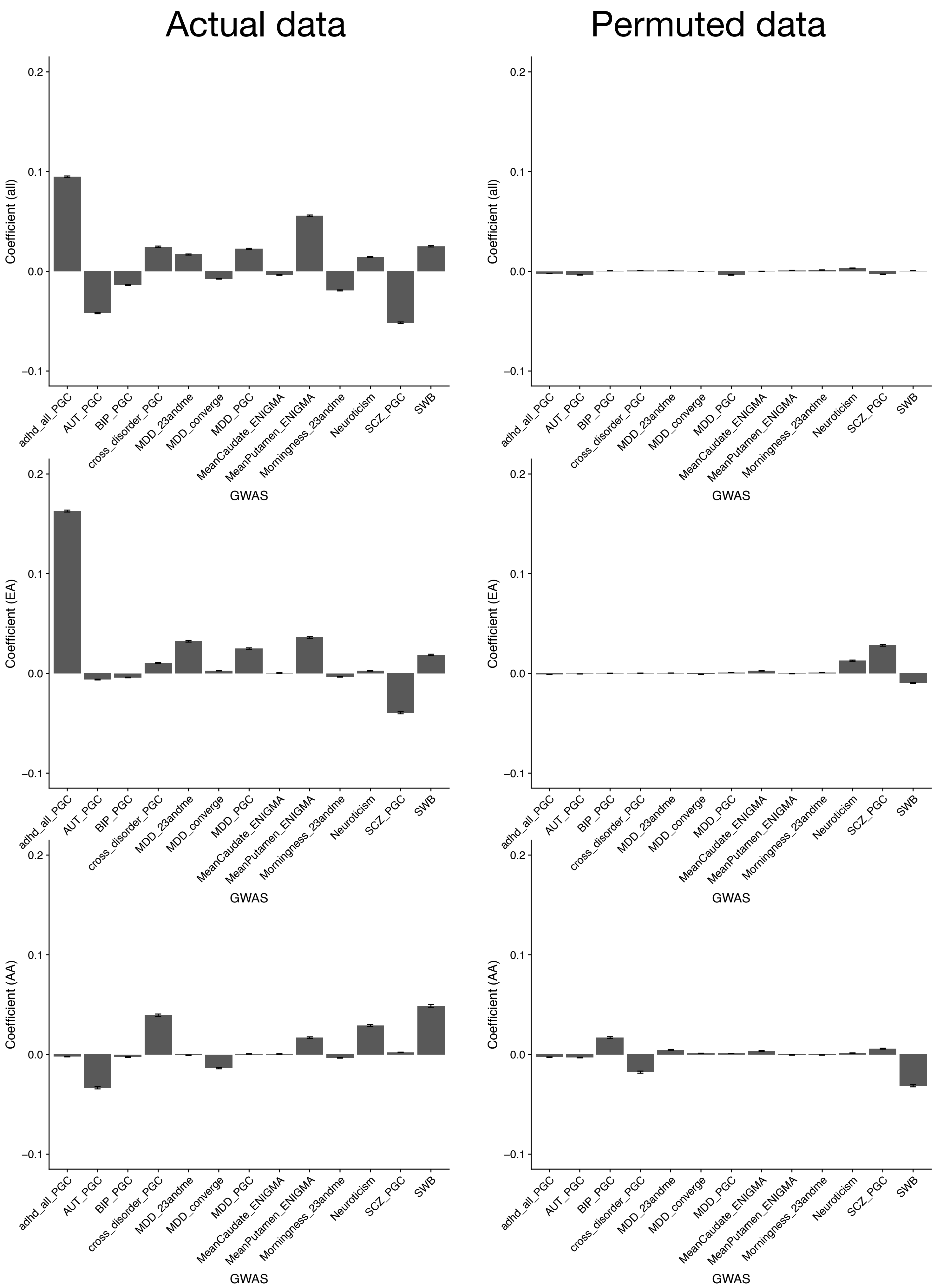

Actual data
Permuted data
Coefficient (all)
Coefficient (EA)
Coefficient (AA)
0.2
0.1
0.0
−0.1
adhd_all_PGC
AUT_PGC
BIP_PGC
cross_disorder_PGC
MDD_23andme
MDD_converge
MDD_PGC
MeanCaudate_ENIGMA
MeanPutamen_ENIGMA
Morningness_23andme
Neuroticism
SCZ_PGC
SWB
GWAS


S Fig 4

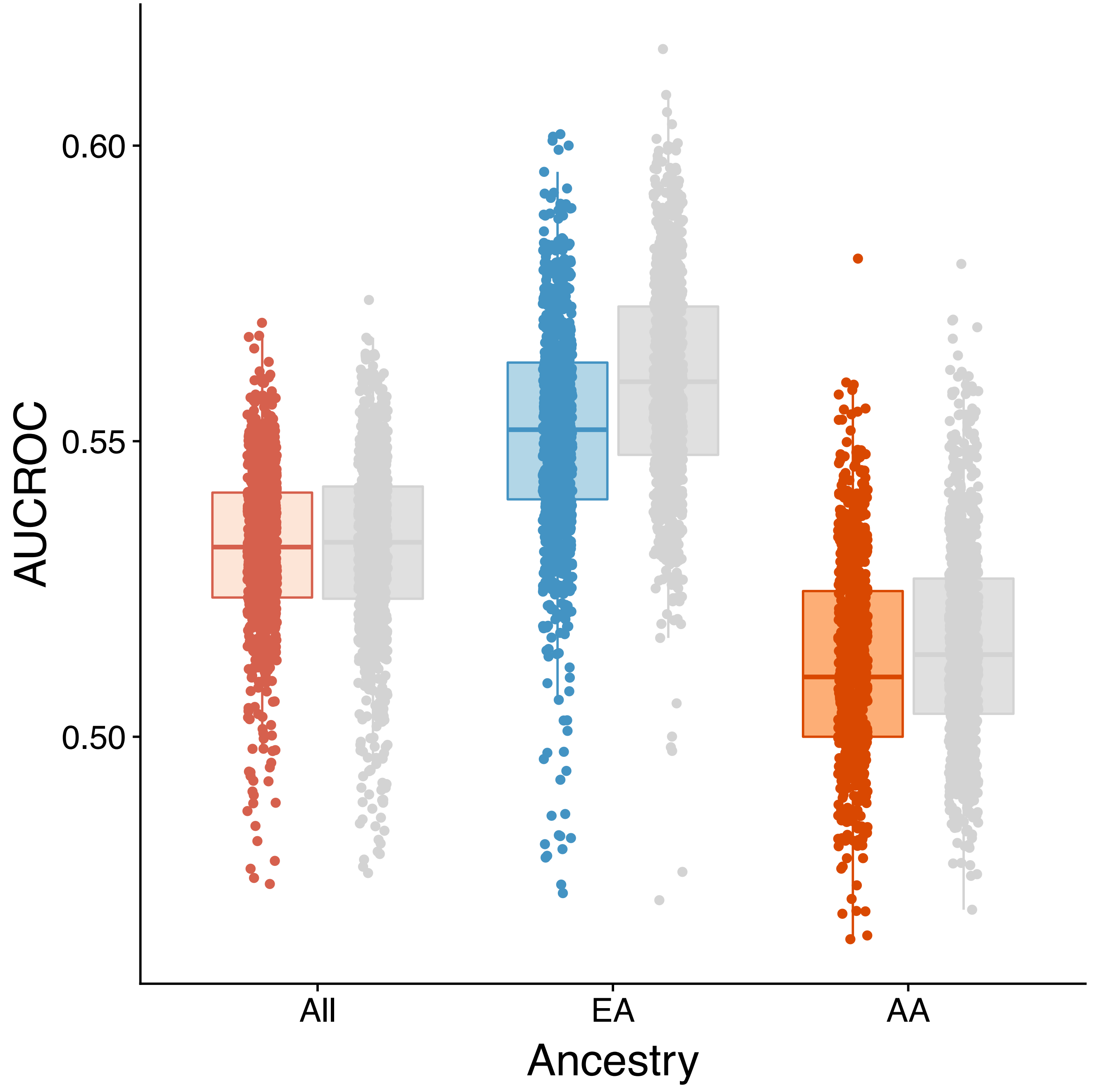


**S Fig 5**

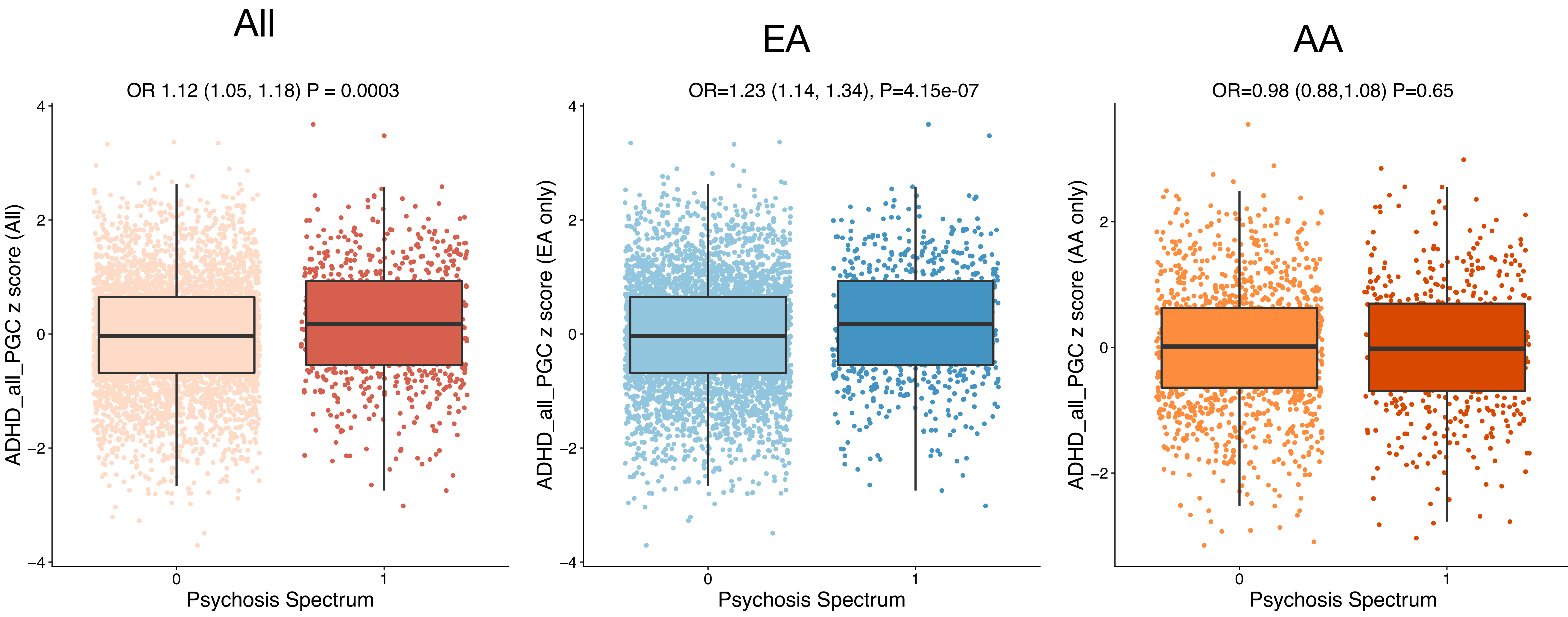

All
OR 1.12 (1.05, 1.18) P = 0.0003
ADHD_all_PGC z score (All)
Psychosis Spectrum
EA
OR=1.23 (1.14, 1.34), P=4.15e-07
ADHD_all_PGC z score (EA only)
Psychosis Spectrum
AA
OR=0.98 (0.88,1.08) P=0.65
ADHD_all_PGC z score (AA only)
Psychosis Spectrum


**S Fig 6**

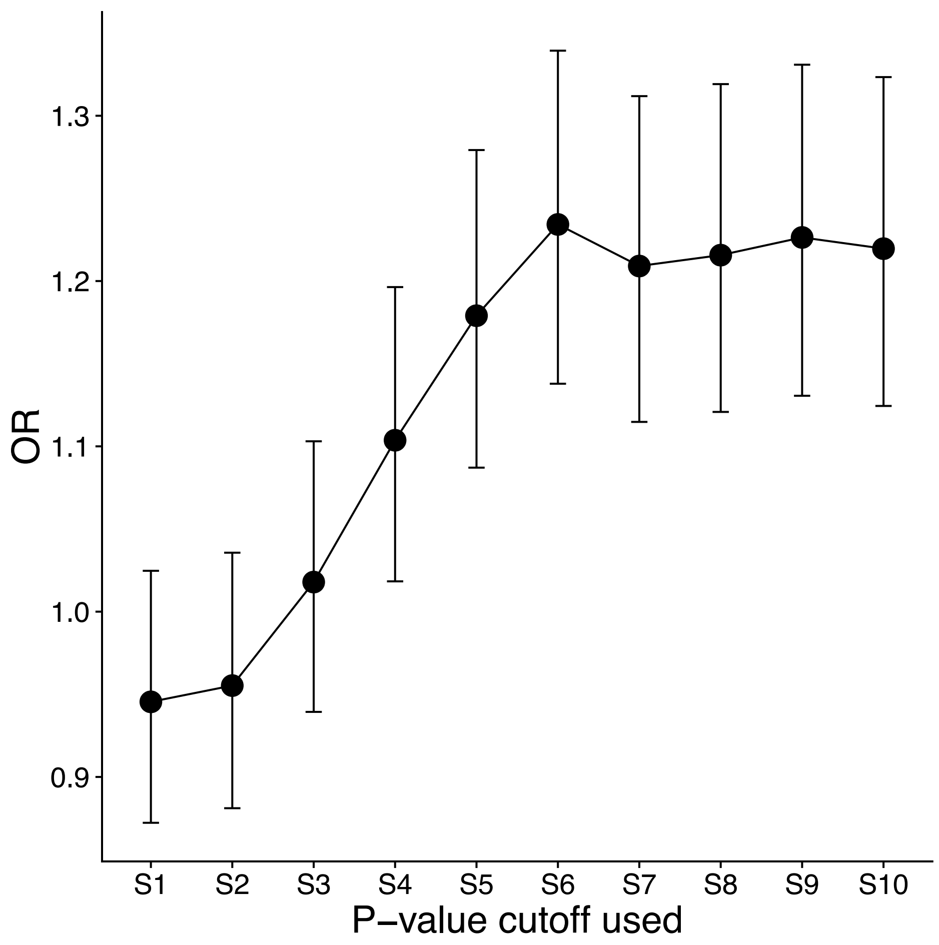


**S Fig 7**

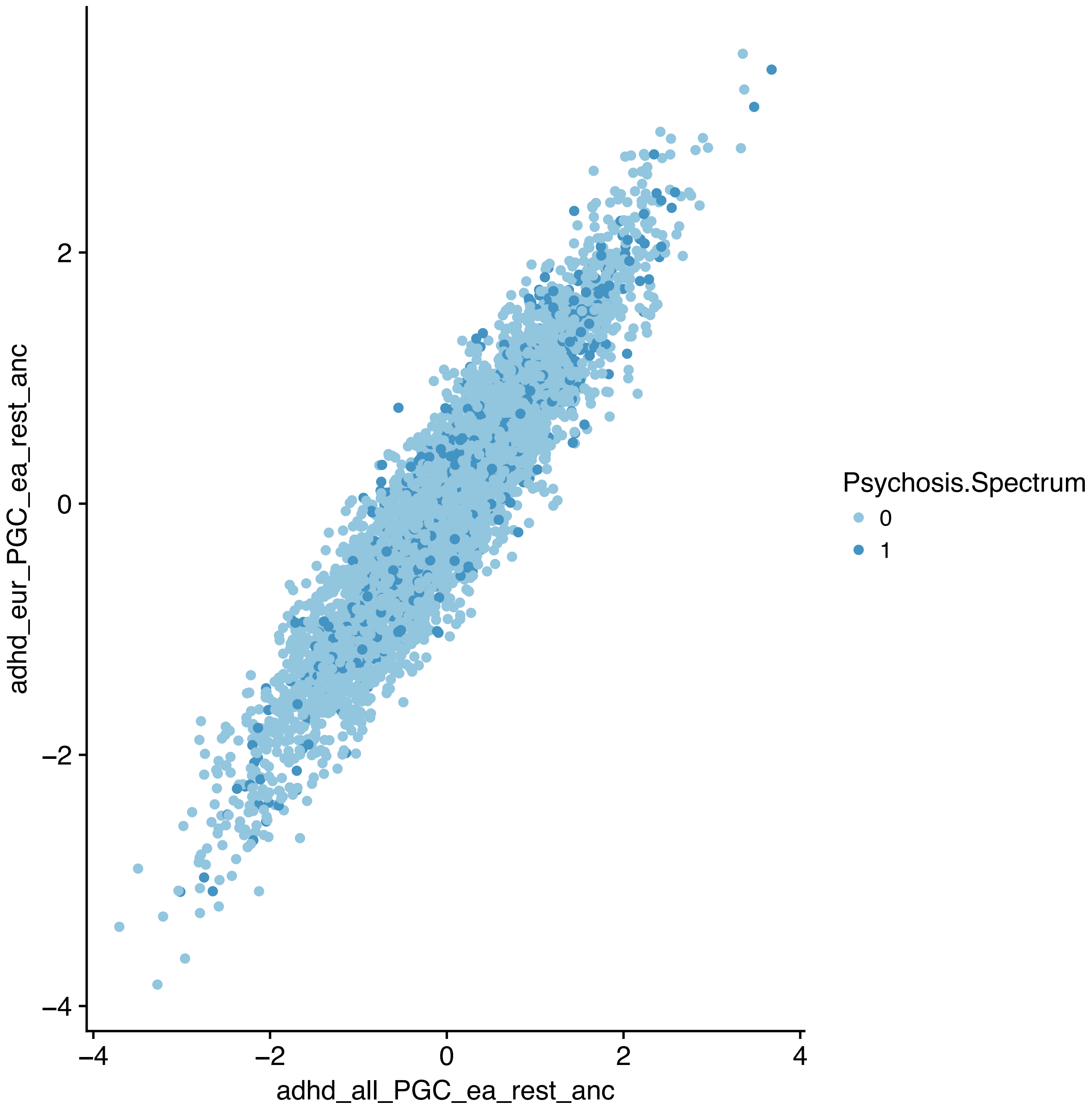

adhd_eur_PGC_ea_rest_anc
2
0
−2
−4
−4
−2
0
2
4
adhd_all_PGC_ea_rest_anc
Psychosis.Spectrum
0
1


**S Fig 8**

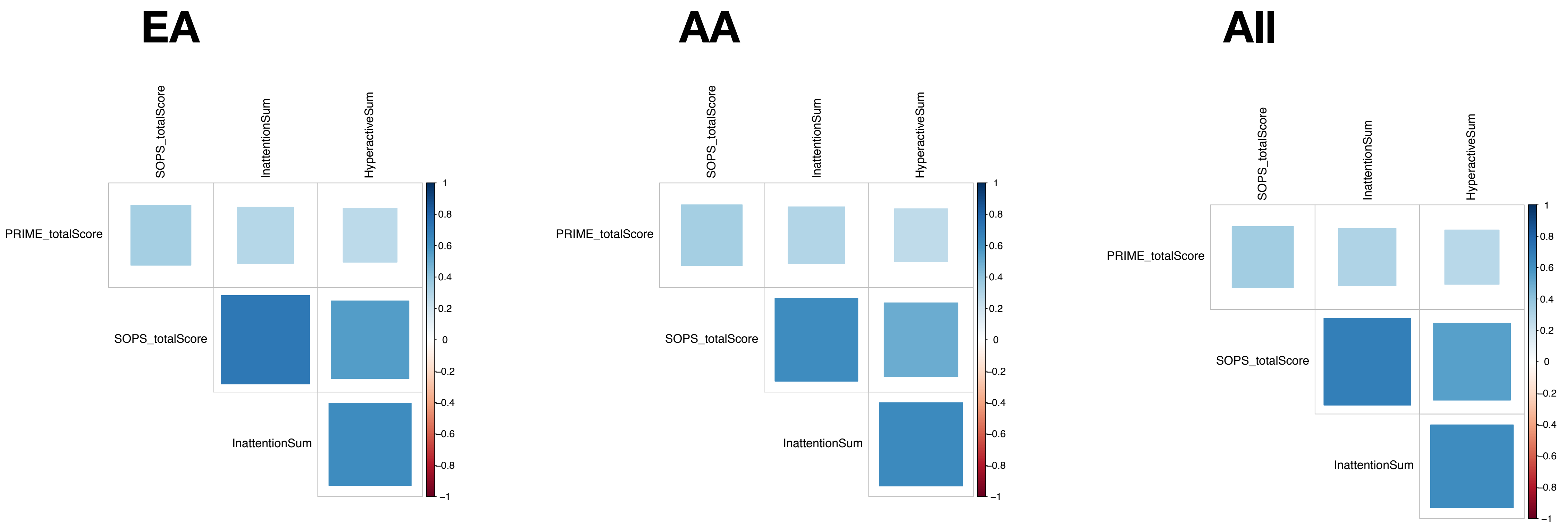
EA
AA
All
SOPS_totalScore
InattentionSum
HyperactiveSum
PRIME_totalScore
SOPS_totalScore
InattentionSum
1
0.8
0.6
0.4
0.2
0
−0.2
−0.4
−0.6
−0.8
−1

S Fig 9

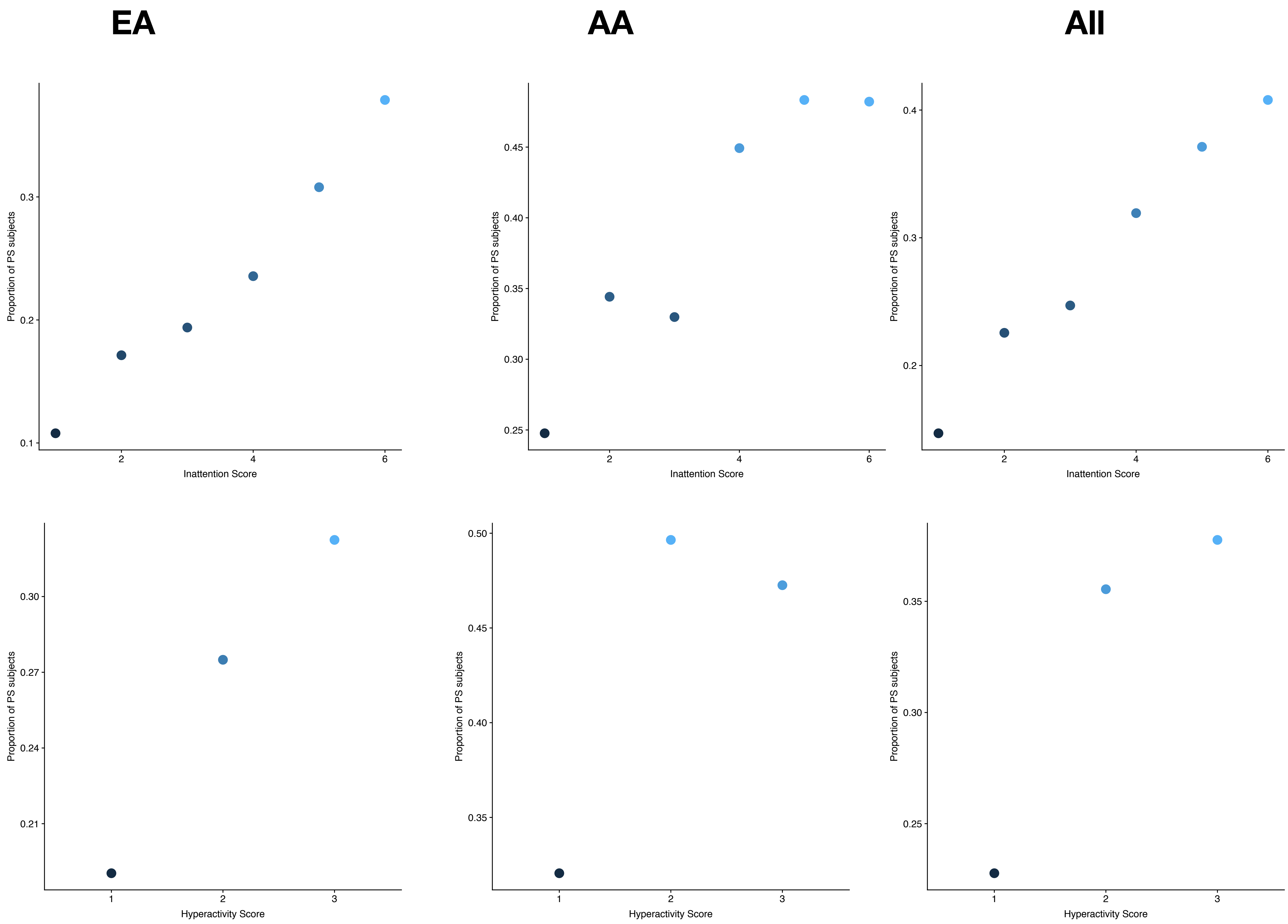

EA
AA
All
Proportion of PS subjects
0.1
0.2
0.3
0.25
0.30
0.35
0.40
0.45
0.4
2
4
6
Inattention Score
0.21
0.24
0.27
0.30
0.35
0.40
0.45
0.50
0.25
0.30
0.35
1
2
3
Hyperactivity Score


S Fig 10

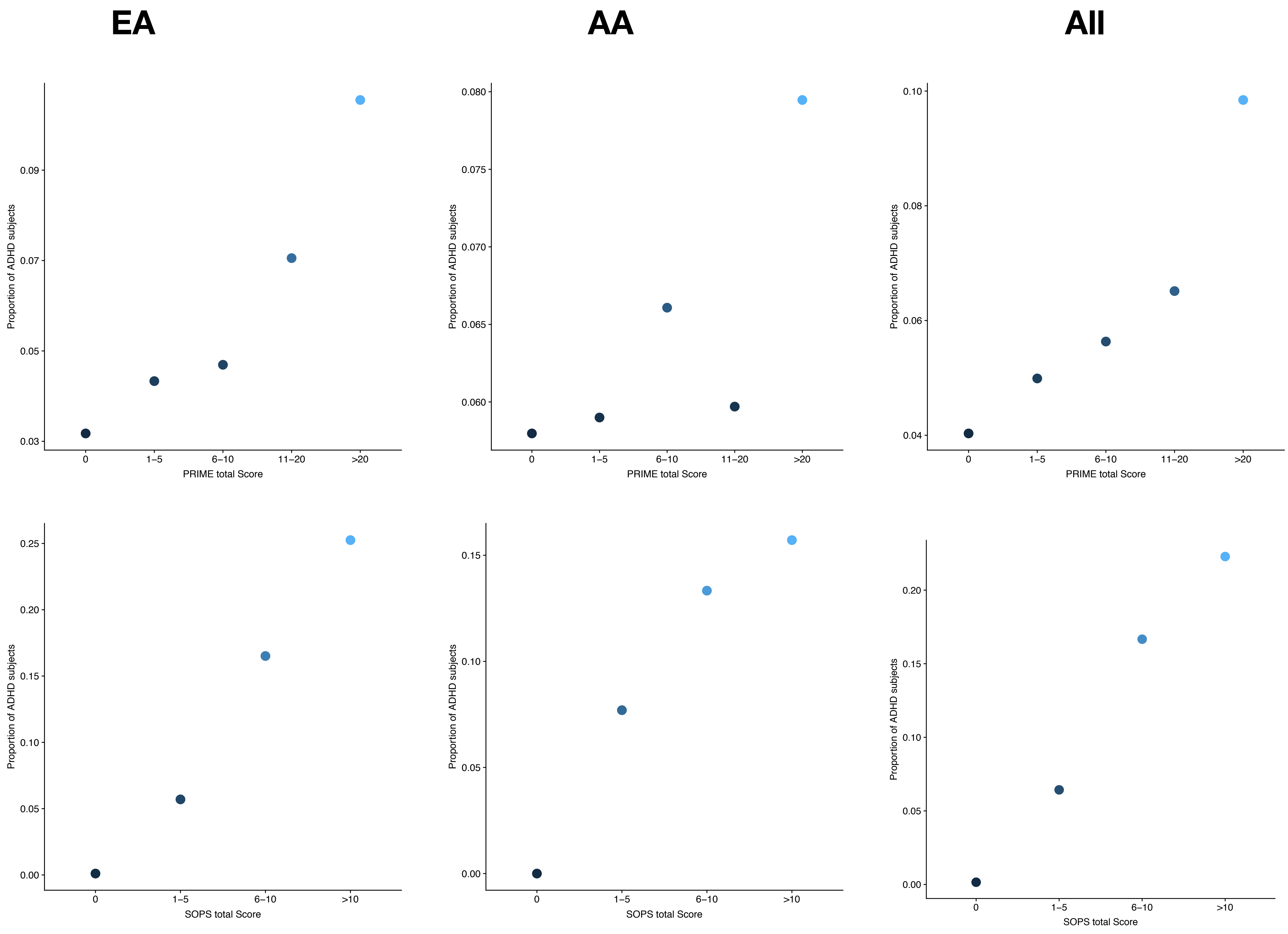
EA
AA
All
Proportion of ADHD subjects
0.03
0.05
0.07
0.09
0.060
0.065
0.070
0.075
0.080
0.04
0.06
0.08
0.10
0
1–5
6–10
11–20
>20
PRIME total Score
0.00
0.05
0.10
0.15
0.20
0.25
>10
SOPS total Score

S Fig 11

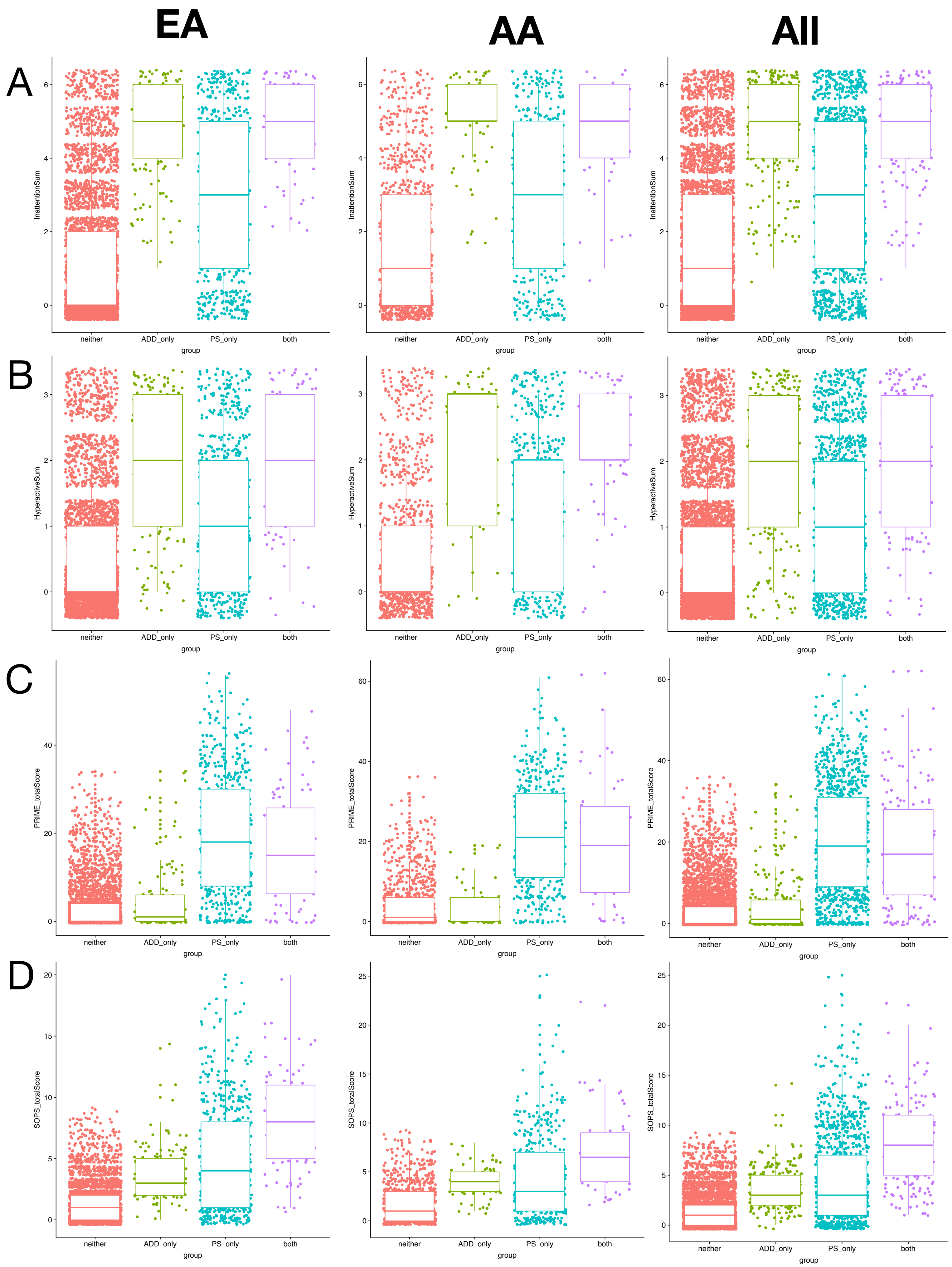
EA
AA
All
A
B
C
D
InattentionSum
HyperactiveSum
PRIME_totalScore
SOPS_totalScore
neither
ADD_only
PS_only
both
group

S Fig 12
